# SARS-CoV-2 maturation driven by a mechano-active nucleocapsid-RNA condensate

**DOI:** 10.64898/2026.09.17.752210

**Authors:** Rini Ravindran, Benjamin Guérin, Ahmad Mahmood, Sherry See Wai Leung, Haytham M. Wahba, Pierre Dagenais, Kaushal Baid, Sauhard Shrivastava, Arkadeb Bhuinya, Sydney Williams, Alisa Sandolache, Pascale Legault, James G. Omichinski, Arinjay Banerjee, Paul W. Wiseman, Adam Hendricks, Stephen Michnick

**Author notes:** Contributed equally (Rini Ravindran, Benjamin Guérin).

## Abstract

The nucleocapsid (N) protein, the core component of the SARS-CoV-2 virus, binds to the 30-kb viral genomic RNA (vgRNA) to form ribonucleoprotein assemblies that are packaged into ∼100 nm membrane-enclosed virions. In addition to viral assembly, the N protein performs other functions, including roles in viral mRNA transcription, replication, and immune regulation, making it a key target for developing diagnostics and vaccines. Recent studies show that N protein and RNA undergo phase separation to form a biomolecular condensate that constitutes the viral core. However, how this condensate becomes selectively enveloped by a membrane remains unclear. Here, using a minimal reconstituted system, we demonstrate that N protein-vgRNA condensates are spontaneously enveloped by lipid bilayer membranes, whereas condensates formed by N protein alone or with genomic RNA fragments fail to undergo envelopment and instead adhere to or weakly deform membranes. We show that RNA length tunes the material properties of N protein condensates, with vgRNA imparting enhanced elasticity. We propose a physical model for SARS-CoV-2 maturation in which adhesion between the membrane and the N protein-vgRNA condensate drives membrane bending, while condensate elasticity preserves core geometry. These results identify RNA-dependent material properties as an underlying physical principle for selective SARS-CoV-2 maturation.

## Introduction

The global impact that the COVID-19 pandemic had on the economy and human health, along with ongoing outbreaks of pandemic potential Middle East respiratory syndrome coronavirus (MERS-CoV) and the detection of multiple pre-emergent bat coronaviruses^1–3^ underscores the need to dissect viral assembly and develop interventions that disrupt the coronavirus life cycle. While extensive research has focused on deciphering the early events of viral entry, the formation of replication-transcription complexes, and immune system responses against the viral load, the critical step of viral assembly remains poorly understood. Coronavirus assembly occurs in the intermembrane space between the endoplasmic reticulum (ER) and the ER-Golgi-intermediate complex (ERGIC)^4^, during which the nucleocapsid (N) protein binds to the 30-kb vgRNA to package the SARS-CoV-2 virion into a 100-nm particle. Although the N protein can bind sub-genomic viral RNA fragments and host RNA^5^, SARS-CoV-2 whole genomic 30-kb RNA is preferentially packaged in purified virus particles^6,7^, likely guided by packaging signals within the vgRNA which contain special stem-loop structures that are thought to govern this specificity^8,9^. How N protein-coated vgRNA interacts with localized areas enriched in membrane (M) protein^10,11^ and the spike (S) protein to form vesicles containing the viral RNA-nucleoprotein (RNP) complexes remains unclear.

SARS-CoV-2 N protein undergoes liquid-liquid phase separation *in vitro*, and multiple *in vivo* functions associated with N protein condensates have been proposed^12–20^. Homotypic N protein interactions are sufficient for phase separation, while heterotypic N protein-vgRNA interactions regulate the efficiency of this process^21^. N protein-RNA condensates likely perform multiple roles during the viral cell cycle, and it is plausible that the membrane-bending step during assembly is driven by these condensates^17^. RNP complexes appear to deform membranes at viral assembly sites, as visualized by cryo-electron microscopy of MHV-infected cells^22,23^. This membrane bending step must be crucial for virion assembly, but how this occurs is unknown. A clue that could explain how the membrane could envelope the nucleocapsid comes from *in vitro* studies of liquid-like droplets’ interactions with lipid vesicles, demonstrating that membrane wetting, tubulation and vesicle budding can mutually transform both condensate and membrane morphologies^24–28^.

Biomolecular condensate-dependent membrane remodeling also regulates several essential cellular processes, including endocytosis^29,30^, genome reorganization^31^, synaptic exocytosis^32^, plant tonoplast budding and nanotube formation^33^, biogenesis of secretory storage granules^34^, endosome membrane bending and scission^30^, autophagy^35^, and organelle biogenesis^36^. General models have been developed to describe how biomolecular condensates contact and remodel materials in living cells, based on balances between the material properties of condensates and their adhesion to surfaces^37,38^. Thermodynamically, surface tension and other interfacial behaviors are determined by the surface energy of the condensate, which reflects how proteins, RNA, ions, and solvent molecules partition between the condensate and the surrounding phase^39^. Equilibrium at the condensate interface ultimately governs their ability to wet, adhere to, and deform membranes.

In this study, we hypothesized that membrane bending during SARS-CoV-2 maturation step is driven by N protein-vgRNA condensate adhesion to the membrane surface. To test this hypothesis, we examined N protein-vgRNA condensate interaction with model membranes and quantified the material properties of the condensates. Our results support a model in which the N protein-vgRNA-condensates drive membrane envelopment through a balance of the condensate elasticity and interfacial adhesion to the membrane, linking the observed phenotypic material properties to molecular determinants of condensate-membrane interactions.

## Results

### N protein-vgRNA condensates can drive spontaneous membrane envelopment

Recombinant full-length SARS-CoV-2 N protein was purified following expression in bacteria; non-specifically bound bacterial nucleic acids were removed (Extended data Fig. 1a-c); and the protein was labeled with Atto 488. The 30kb SARS-CoV-2 vgRNA was isolated from infected mammalian cells, while smaller viral RNA fragments (1-1.5 kB) were purified by in vitro transcription (Extended data Fig. 1d,e). N protein-Atto 488 phase separated into condensates under physiological buffer conditions both in the absence and presence of vgRNA. Although N protein (2 µM) alone formed condensates, inclusion of increasing vgRNA concentrations enhanced phase separation, resulting in larger condensates with variable sizes (Fig. 1a-c). These observations are consistent with RNA length acting as an important determinant of condensate material properties.

**Fig.1:**
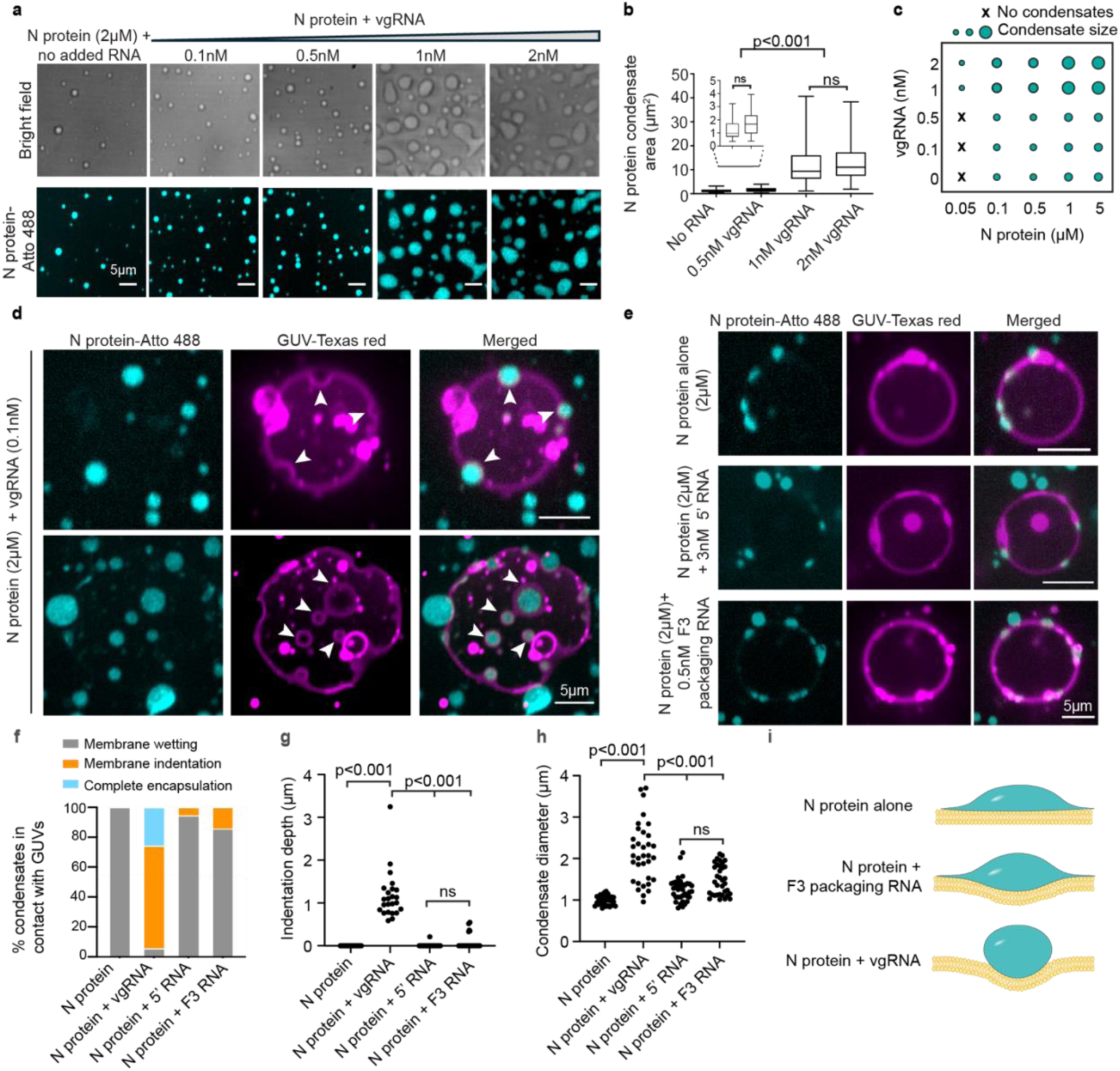
N protein-vgRNA condensates drive spontaneous membrane envelopment. **a**, Bright-field (top) and fluorescence (bottom) images of N protein (2 µM; Atto-488) in the absence or presence of increasing concentrations of vgRNA (0.1–2 nM), which demonstrate that vgRNA promotes the formation of larger condensates. Scale bars, 5 µm. **b,** Condensate area of N protein as a function of vgRNA concentration. Boxes show median and interquartile range; whiskers indicate minimum and maximum values. **c,** Phase diagram of N-protein-vgRNA. **d,** Confocal images of N-protein condensates (cyan) in the presence of vgRNA (0.1 nM) interacting with giant unilamellar vesicles (GUVs; Texas Red, magenta), showing membrane bending events (top) complete encapsulation of condensates (bottom). White arrowheads indicate condensates associated with GUVs. Scale bars, 5 µm. **e,** N protein condensates wet the GUV membrane in the absence of any added RNA (top), 5′ end RNA (center) and packaging RNA (bottom), but do not display membrane encapsulation or significant bending. **f,** Quantification of N protein condensate interaction with GUVs observed under the conditions shown in **d** and **e**. **g,** Extent of GUV membrane indentation induced by N-protein condensates. **h,** N-protein condensate size under the indicated conditions. **i,** Model illustrating RNA-dependent N protein condensate interaction at membranes and associated membrane deformation.

To test the hypothesis that N protein-vgRNA condensates are sufficient to drive membrane deformation, we added giant unilamellar vesicles (GUVs) to freshly formed N protein condensates and assessed condensate-membrane interactions. N protein-vgRNA condensates retained their spherical shape upon contact with GUVs and induced localized membrane invaginations (Fig. 1d top), as well as fully membrane-encapsulated, condensate-filled structures within the GUVs. (Fig. 1d bottom). Similar behavior has also been reported for the plant ESCRT component FREE1, which forms condensates that can bend and induce fission in GUVs independently of other ESCRT component^40^. In contrast, the N protein condensates formed in the absence of viral RNA “wetted” the GUV surface without membrane deformation (Fig. 1e top). Comparable membrane-wetting interactions were observed for N protein condensates formed with a 1-kb 5′ vgRNA fragment (Fig. 1e middle), previously shown to enhance phase separation of N protein in vitro^21^. At higher 5′RNA concentrations, condensates retain a spherical shape on contact with GUVs (Extended data Fig. 2a).

We also examined N protein condensates formed with a 1.4 kb packaging signal RNA (F3 sequence) spanning the Nsp12–Nsp13 coding regions, previously reported to promote efficient RNA packaging^9^ (Fig. 1e bottom). Unlike the 5′ RNA fragment, the F3 packaging RNA induced localized membrane bending at higher RNA concentrations (Extended data Fig. 2b), suggesting that sequence-encoded features can contribute to membrane remodeling even at shorter RNA length scales. However, the extensive membrane invaginations and fully encapsulated morphologies observed with full-length vgRNA were not reproduced by either of the shorter viral RNAs tested (Fig. 1f,g; Extended data Fig. 2c,d). Together, these findings suggest that RNA length is a dominant determinant of condensate-driven membrane remodeling, while sequence-dependent interactions can further modulate this behavior.

Images were analyzed without selection bias, as condensate–GUV interactions occurred stochastically and could not be experimentally controlled. All interacting GUVs, across a broad size range, were included (Extended data Fig. 2e). Furthermore, relative deformation, quantified as percent indentation (Indentation depth/Condensate diameter), showed no significant dependence on GUV diameter (linear regression: R^2^= 0.02345 p= 0.4192; Spearman correlation r= 0.08478 p= 0.6560 (Extended data Fig. 2f)). These results indicate that variability in GUV size is unlikely to be a major determinant of the observed RNA-dependent condensate–membrane interactions.

We note that N protein-vgRNA condensates were significantly larger than condensates formed by N protein alone or with shorter viral RNA fragments (Fig. 1h). The range of N protein and RNA concentrations tested were guided by previous studies^21^ and titrations optimized for spherical condensate formation (Fig.1c). At packaging RNA concentrations above 0.5 nM, N protein formed elongated and aggregate-like condensates (Extended data Fig. 2g,h) that were unsuitable for GUV assays. This could be due to the multivalent interactions among stem-loop structures present in this region of the SARS-CoV-2 genome^41^ that could change and influence phase separation. Although the packaging sequence is present in the vgRNA, higher-order secondary structures and long-range RNA interactions may modulate its accessibility and behavior. Overall, these observations support a model in which RNA-dependent modulation of condensate material properties, strongly influenced by RNA length, enables membrane deformation and envelopment. Within this framework, the SARS-CoV-2 genomic RNA represents a biologically relevant example of a long, multivalent RNA whose sequence-encoded features may further enhance membrane remodeling (schematized in Fig. 1i).

### vgRNA imparts increased elasticity and cohesion to N protein-vgRNA condensates

Given that condensate–GUV interactions differ depending on the RNA species present in the condensate (Fig. 1), we tested whether the identity of the RNA alters the surface charge of the condensates, which could in turn modulate condensate–membrane adhesion. Zeta potential measurements revealed that N protein condensates are weakly positively charged (+3 to +5 mV), and that at the low RNA concentrations used in GUV experiments they do not significantly alter this value, regardless of the identity of the RNA (Extended data Fig. 3a). Only at higher concentrations did excess negative charge from the RNA shift the zeta potential to negative values (Extended data Fig. 3b). Thus, the distinct membrane interaction behaviors observed for condensates formed with different RNA species cannot be explained by differences in net charge, implicating RNA-dependent changes in condensate material properties as the dominant factor.

Next, we measured the effect of RNA on the viscoelasticity of N protein condensates using optical tweezers. *In vitro*, the N protein condensates are typically 1 – 2 µm in radius, making it difficult to internalize probe beads into the condensate to perform conventional optical tweezer micro-rheology^29^. Hence, we used an optical tweezer force indentation assay to probe the surface mechanics of N protein-RNA condensates (Fig. 2a left-middle). We find that for indentations near the condensate’s surface; condensates exhibit a primarily elastic response (Extended data Fig. 6e). Following previous studies, we modeled optical trap-driven soft matter interactions using idealized Hertzian contact models^42,43^. Measuring soft matter deformations at sub-micron scales is challenging due to unaccounted for probe-sample interactions and the inherent length scales of the sample^44^. Thus, in practice, we assume linear elasticity at small strains when applying Hertz theory and Hertzian contact models^44^. The force signal, measured using back-focal plane interferometry, was plotted as a function of probe depth to produce force-indentation curves (FICs) (Fig. 2a, right). The initial contact region [0 nm – 200 nm] of the ensemble average FICs (Fig. 2b) were fitted to a Hertzian contact model to estimate the condensate’s apparent modulus (Fig. 2c) which is a combination of viscous, elastic, and bulk contributions imparted by the condensate’s interface^42^. Condensate fusion experiments reveal that while RNA content mediates the condensate’s viscosity to surface tension ratio (Extended data Fig. 4) – condensate fusion events result in complete fusion with spherical equilibrium geometries irrespective of RNA content. Previous experimental and computational modeling of condensate fusion revealed that condensate composition, internal organization, and aging alter bulk condensate viscoelasticity which in turn leads to asymmetric fusion events^45^. We do not observe asymmetric condensate fusion suggesting that while RNA content may mediate internal rearrangement time scales via transient homotypic and heterotypic interactions, N protein condensates remain predominantly liquid like wherein long-time scale elasticity is primarily imparted by surface tension.

**Fig.2:**
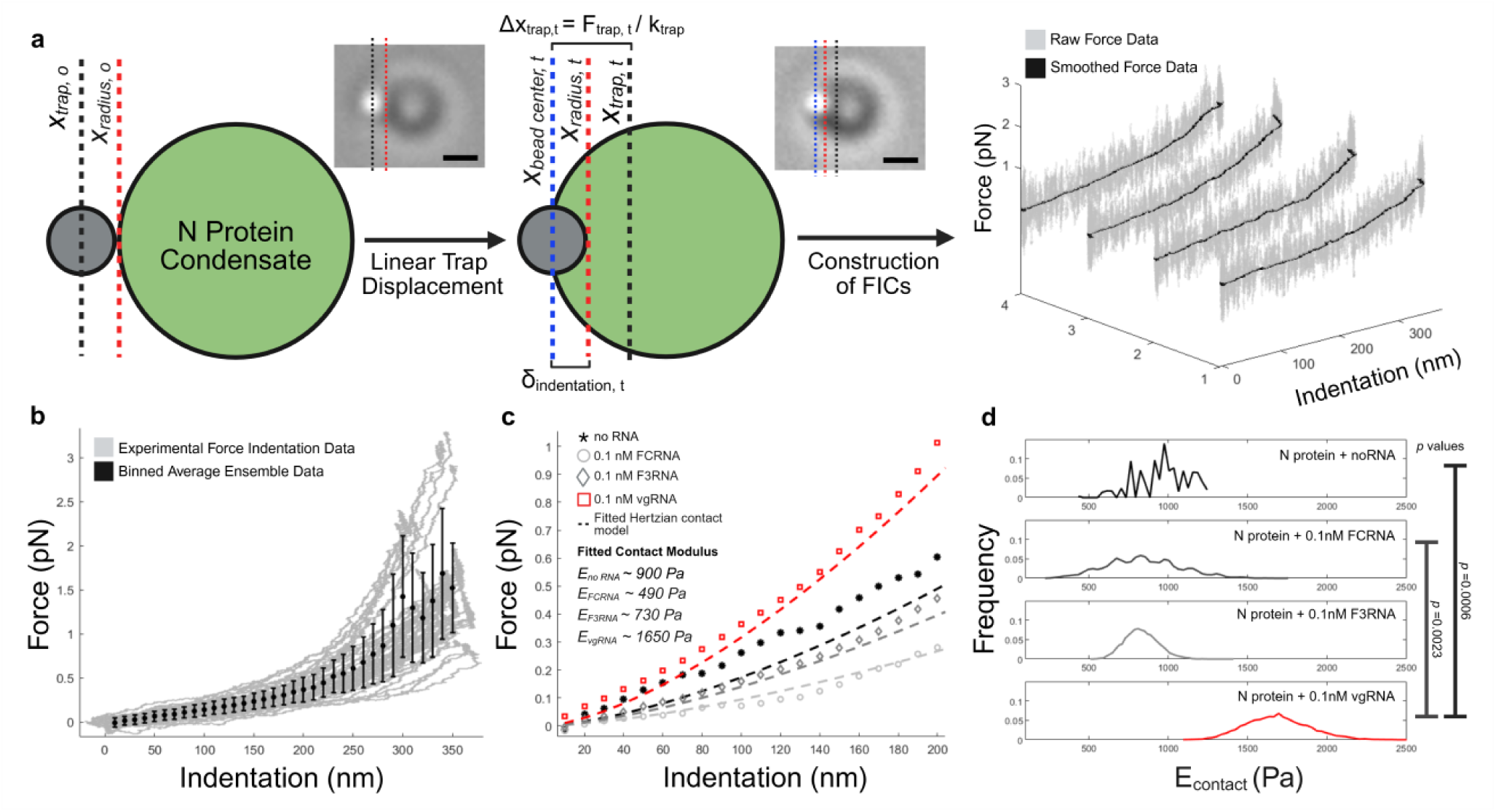
vgRNA increases stiffness of N protein-vgRNA condensates. **a,** Schematic and brightfield image (inset) of the optical trapping indentation assay. Position of trapped probe relative to condensate before (left) and after (middle) linear trap displacement. Scale bar, 1 µm. **b,** All force indentation curves (grey) for 1 experiment of the vgRNA condition. Black symbols indicate the ensemble average and standard deviation. **c,** Ensemble average force indentation curves (symbols) fitted to a Hertzian contact model (dashed lines). The Hertzian contact model fit was used to estimate the ensemble elastic modulus (E_vgRNA_ ∼ 1650 Pa; E_F3RNA_ ∼ 730 Pa; E_FCRNA_ ∼ 490 Pa; E_no RNA_ ∼ 900 Pa). **d,** We applied the Hertzian contact model to individual force indentation curves to estimate the average Young’s modulus of each condensate (N_vgRNA_= 31 condensates; N_F3RNA_= 19 condensates; N_FCRNA_= 11 condensates; N_noRNA_= 4 condensates). We then obtained the average distribution from bootstrapped (N_bootstrap_=10000) samples. *P* values were calculated from the difference of bootstrap means.

The apparent modulus for N protein-vgRNA (∼1650 Pa) was approximately two-fold higher than N protein alone condensates (∼900 Pa) (Fig. 2c,d). Unlike the vgRNA, the packaging sequence RNA (F3 RNA) and a size control for the packaging RNA (FC RNA) did not increase the apparent moduli of the N protein-RNA condensates (Fig. 2c,d). Indentation-retraction curves appear highly elastic with no observable probe-condensate adhesion, suggesting that we are probing the N protein condensate’s interfacial elasticity dominated by its interfacial tension (Extended data Fig. 5-6e). If we interpret the apparent moduli as measures of the Laplace pressure across the N protein condensate dense-dilute phase interface, we obtain interfacial tension estimates that are on the order of mNm^−1^. This order of magnitude is consistent with micro-pipette aspiration measurements performed on membrane bending condensates^27^. At large indentation depths, we observe deviations from the idealized Hertzian contact model where force is expected to increase exponentially with respect to indentation depth (*F(d) ∝ d*^3^*^/^*^2^) (Extended data Fig. 6a-b). While all RNA conditions showed a similar deviation at indentation depths that exceeded the probe radius (Extended data Fig. 4b-c), vgRNA-N protein condensates showed additional deviation for lesser indentation depths. Taken together, these data suggest that the RNA content within N protein condensates modulates its interfacial elasticity and imparts additional non-linear and strain stiffening effects with increasing or repeated strain. In the context of SARS-CoV-2 viral particle maturation, such enhanced elasticity may enable N protein condensates to more effectively deform and remodel cellular membranes.

### N protein and RNA within condensates show nanoscale inhomogeneities

The increased stiffness and non-linear mechanical response of N protein-vgRNA condensates (Fig.2) suggest that their emergent material properties arise from internal dynamical molecular organization spanning different length and time scales^46^. To investigate sub-micrometer molecular behavior on microsecond time scales, we performed fluorescence lifetime correlation spectroscopy (FLCS). To probe micrometre-scale molecular reorganization within the condensate over sub-second timescales, we performed fluorescence recovery after photobleaching (FRAP), and spatiotemporal image correlation spectroscopy (STICS).

To determine the spatial organization of N protein and RNA within the condensates, RNA was labeled with Cy3 at the 3’ hydroxyl end and N protein with AF488 via maleimide-thiol chemistry on a C-terminal Cys residue inserted in the sequence at the C-terminus, ensuring one dye per molecule for both the N protein and vgRNA. The N protein-AF488-vgRNA-Cy3 condensates were imaged using a time-correlated single-photon counting confocal microscope revealed distinct dense, dynamic fluorescence-rich nodes (Fig.3a). Similar heterogeneity was observed for vgRNA-Cy3 partitioned within the condensates (Fig.3b). N protein and vgRNA nodes partially overlap with a size range of 270 ± 30nm (Fig. 3c) and exhibit correlated movement over time, suggesting coordinated reorganization of protein and RNA within the condensate interior. To quantify the dynamics of intra-condensate clusters, we applied a wavelet-transform-assisted STICS approach^47,48^. A Richter wavelet filter tuned to the size scale of the observed clusters accentuated mesoscale features, enabling the extraction of average cluster velocities. This analysis revealed slow collective movements of both N protein and RNA clusters within the condensate (Extended data Fig. 5a-f, Supplementary videos 1-3). Such spatial inhomogeneities have been reported in coacervate condensates composed of oppositely charged proteins^49^, and are predicted to influence molecular dynamics in nucleolar granular compartments^50^. Similarly, condensates formed by prion-like low-complexity domains exhibit slow-moving nanoscale hubs stabilized by reversible aromatic interactions^51^. In N protein-vgRNA condensates, the observed nanoscale hubs likely reflect charge partitioning within intrinsically disordered regions (IDRs) of the N protein and the formation of higher-order assemblies, contributing to elastic resistance while preserving internal rearrangement.

**Fig.3:**
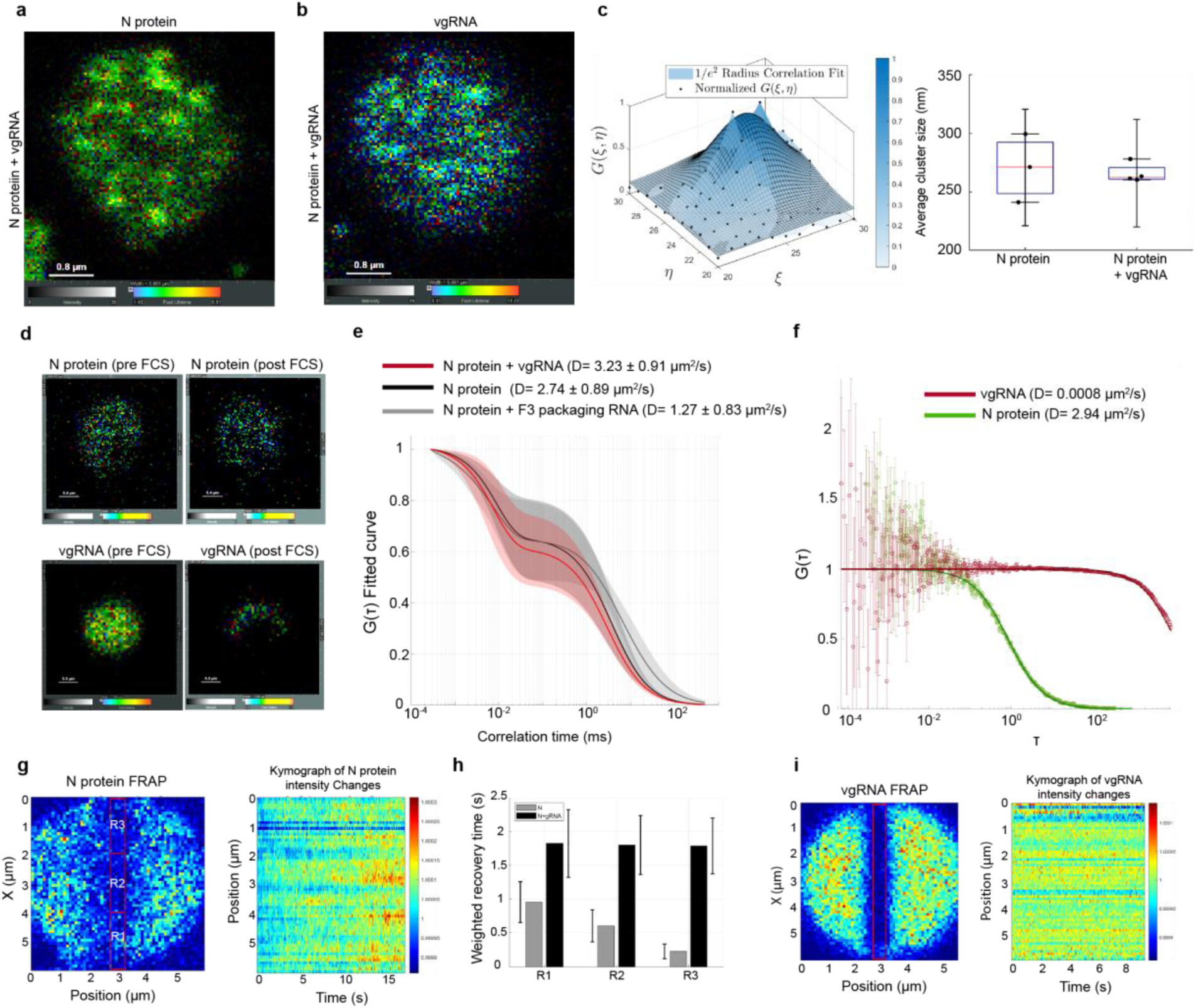
N protein and vgRNA form dynamic, fluorescence-rich clusters within condensates. **a,** N protein-AF488 and **b,** vgRNA-Cy3 image of N protein-vgRNA condensates at low labelling concentrations imaged with a TCSPC microscope. Images are weighted by the fluorescence lifetime. **c,** An imaging time series of condensates was analyzed frame by frame with image correlation spectroscopy (ICS) to extract the average length scale of intra-condensate clusters. Normalized spatial autocorrelation functions (ACF) were fitted with an asymmetric Gaussian function. Fit parameters *ω_x_* and *ω_y_* (*e*^−2^ radius) were averaged over 128 frames for each condensate (3 in the presence of vgRNA, and 4 without). **d,** Fluorescence lifetime correlation spectroscopy (FLCS) measurements of N protein-AF488 were taken inside condensates formed in the presence of vgRNA and packaging RNA. Images were taken prior to and after the 120 s measurement time. **e,** Lifetime filtering for ACF calculation was performed by fitting the lifetime histogram to a 3 exponential decay and selecting for the 4.1 ns lifetime of AF488. FLCS ACF curves were fitted to a single diffusing species with a triplet state. **f,** A representative FLCS curve for an N protein-vgRNA condensate displaying the multiple decade separation in mobility. **g,** Representative image of N protein-AF488 condensate post centered confocal line scan of length 5 µm with 2000 repetitions of 488 nm laser line at 1 mW. Intensities at regions R1, 2, and 3, were analyzed separately for average intensity recovery over time. A kymograph representation of the normalized intensity recovery in time along the length of the condensate, averaged over a width of 5 pixels**. h,** Normalized intensity recovery curves of N protein-AF488 were fit with a simple one-step exponential rate to extract a recovery time in regions R1, 2 and 3 for condensates formed in the presence and without vgRNA. A weighted average was performed for a sample of N=4**. i.** Representative image of N protein-AF488 condensate formed in the presence of vgRNA-Cy3 post-centered confocal line scan of length 5 μm with 2000 repetitions of 561 nm laser line at 200 µW with a kymograph representation of the normalized intensity recovery in time along the length of the condensate, averaged over a width of 5 pixels.

We next measured the diffusion of N protein and RNA in individual condensates using FLCS (Fig.3d). By incorporating fluorescence lifetime information, FLCS improves discrimination between overlapping signals or emission of a probe in different molecular environments, enhancing specificity relative to conventional fluorescence correlation spectroscopy. The diffusion coefficient of N protein in the presence of vgRNA is marginally higher than for N protein alone(3.23 ± 0.91 and 2.74 ± 0.89 µm^2^/s, respectively) (Fig.3e), although variability between condensates rendered the difference non-significant. In contrast, N protein condensates formed with F3 packaging signal RNA exhibit a lower protein diffusion coefficient (1.27 ± 0.83 µm^2^/s). In conventional dilute solutions, diffusion coefficients are inversely related to viscosity through the Stokes–Einstein relation. In biomolecular condensates, however, molecular motion often cannot be described as simple Brownian diffusion in a homogeneous Newtonian fluid. Multivalent interactions can generate transient crosslinks and condensate-spanning networks, producing viscoelastic behavior, spatial heterogeneity, and time-dependent changes in material state^52–54^.

Consistent with this, FRAP measurements show that N protein fluorescence recovers only partially (50-80%) following bleaching in both N protein-vgRNA and N protein alone condensates, indicating the coexistence of mobile and immobile protein fractions (Extended data Fig. 6a,b). Line-FRAP measurements with higher temporal resolution revealed slower recovery of N protein in the presence of vgRNA (Fig. 3g,h), suggesting increased connectivity within the condensate network. Conversely, vgRNA exhibits negligible recovery in both FLCS and FRAP measurements (Fig.3f,i, Extended data Fig. 6c), indicating local immobilization on short length and time scales. However, vgRNA displayed collective mobility at longer length scales, as revealed by STICS analysis (Extended data Fig. 5e,f), demonstrating that vgRNA is locally constrained yet capable of coordinated motion within the condensate. The correlated motion of positively charged N protein clusters and negatively charged vgRNA clusters suggests a spatially heterogeneous electrostatic network. Such charge-frustrated architectures are characteristic of polyelectrolyte systems and may contribute to the nonlinear force–indentation responses observed in mechanical measurements (Fig.2), as local rearrangements of charged domains resist deformation while allowing collective reorganization.

A marginally higher diffusion rate of N protein in the presence of vgRNA, despite increased bulk stiffness, indicates that enhanced elasticity arises from increased network connectivity mediated by transient interactions rather than simple viscous slowing. Dynamic N protein-RNA binding, proposed to facilitate scanning of the viral genome for high-affinity sites^55^ is compatible with rapid local diffusion of N protein within a mechanistically cohesive network, whereas the F3 packaging signal RNA, likely promotes more stable binding conformations that reduce protein mobility (Fig.2c, 3e). Although binding kinetics cannot be directly resolved here, these interpretations are supported by consistent trends across FLCS, FRAP, and STICS measurements. Diffusion measurements were performed using picomolar concentrations of fluorescently labeled N protein to minimize perturbation of condensate architecture. Higher labeling stoichiometries enhanced fluorescence signals but also promoted imaging-induced crosslinking (Extended data Fig. 6d,e), constraining absolute diffusion estimates while preserving relative comparisons across conditions. Together, these results indicate that vgRNA promotes a cohesive yet dynamic internal architecture that combines enhanced connectivity with retained molecular mobility, consistent with the elevated Young’s modulus and membrane-deforming capacity of N protein–vgRNA condensates.

### Mechanical properties of N protein-vgRNA condensate determine GUV membrane bending

N protein condensates display a range of membrane interactions depending on the protein’s phosphorylation state and the presence of RNA^56^. We observe that N-protein-vgRNA condensates selectively induce membrane bending and encapsulation (Fig.1), highlighting the specificity of RNA-dependent membrane interactions. Optical trap measurements revealed that RNA alters condensate mechanics and interfacial elasticity (Fig.2), while high-resolution microscopy uncovered nanoscale heterogeneities and differential mobilities of RNA and protein (Fig.3), indicating a relatively immobile RNA scaffold embedded within a dynamic, network-forming protein matrix. Together, these observations suggest that condensate geometry, viscoelasticity, and internal heterogeneity jointly influence how condensates interact with membranes. To quantitatively connect these features, we consider a free energy model incorporating interfacial, contact line, and membrane bending energies^24,25,27,35^ (see methods). Using condensate–membrane images, we extracted intrinsic contact angles and maximum membrane deflections and identified stable energy landscape solutions to estimate relative energy scales for membrane bending.The intrinsic contact angle (θ_C_) of the condensate with the GUV membrane is determined by the mechanics of the N protein RNA co-condensate and the membrane^56^. This intrinsic contact angle is maintained for different membrane deflection geometries. Assuming that N protein condensate volume is constant on the timescale of the membrane interaction, the N protein condensates adopt a double spherical cap geometry where θ_C_ is conserved across a range of above (*φ*) and below (*φ*) membrane deflection angles (Fig. 4a-b; Extended data Fig. 9). N protein condensates transition to different membrane deflection geometries to minimize their free energy (Extended data Fig. 10a). We measured intrinsic contact angles and above membrane deflection angles from images of condensate-membrane interactions (Fig. 4c) and determined the stable solution of the energy landscapes that best fit the measurements. From these values we could extract estimates for relative energy scales of membrane bending (*κ*_bend_) and three-phase contact line (λL_123_) energy (Fig. 4c; Extended data Fig. 10b). RNA-protein interactions alter the mechanics of the N protein-RNA condensates (Fig. 2), and in turn their interactions with membranes. Through fitting the model to the observed condensate-membrane geometries, we find that in the absence of any RNA, membrane bending energy dominates over the condensate’s interfacial energy (no RNA: *κ*_bend_/*γ*_DS_R^2^ = 6.07) (Fig. 4c). Under these conditions, condensates deform to wet the membrane. In contrast, condensate interfacial energy increasingly dominates in the presence of RNA (F3RNA: *κ*_bend_/*γ*_DS_R_D_^2^ = 0.55; vgRNA: *κ*_bend_ /*γ*_DS_R_D_^2^ < 0.0001) to drive membrane bending while condensates remain nearly spherical.

**Fig.4:**
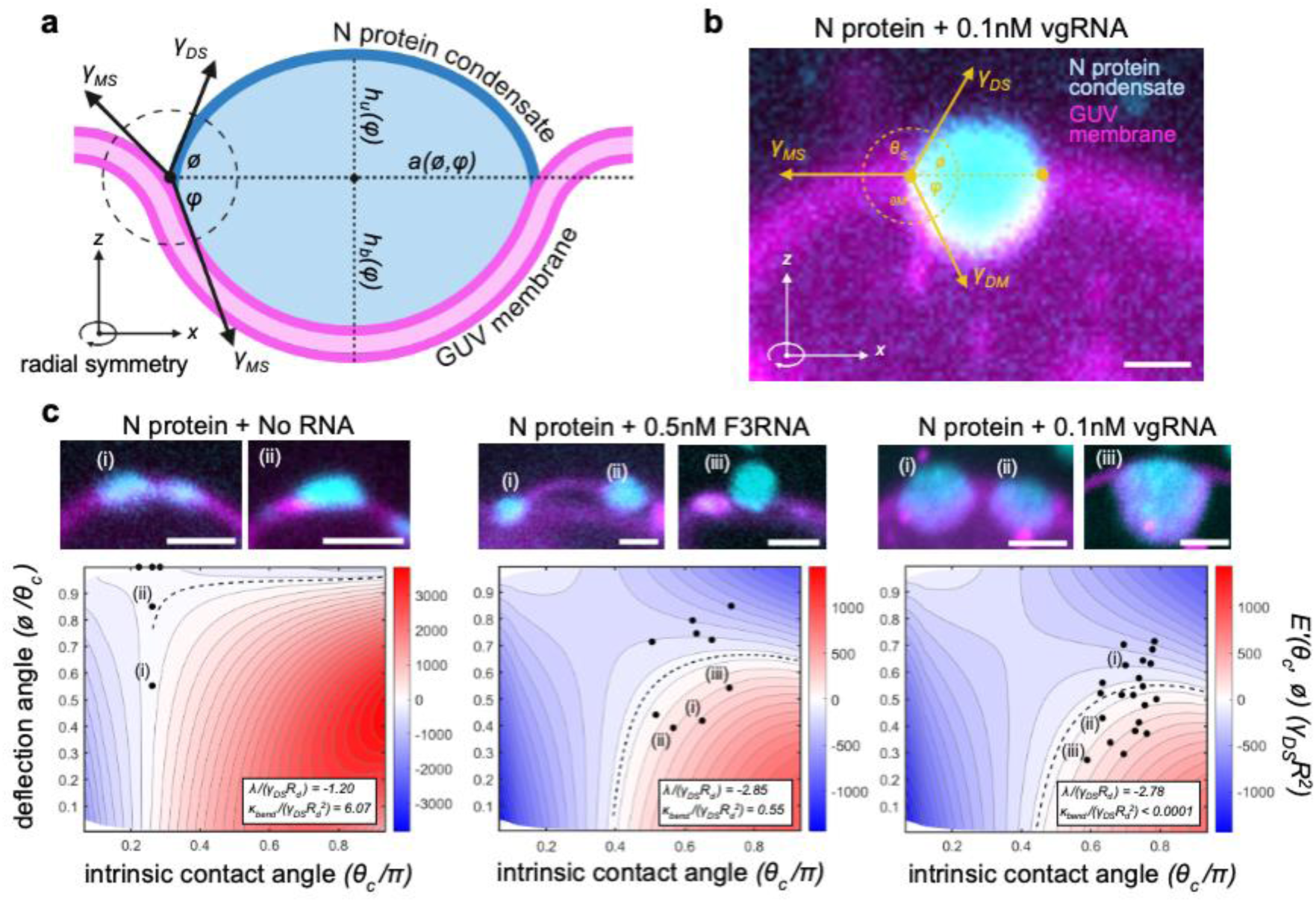
N protein condensates differentially interact with GUV membranes depending on RNA content. **a,** Schematic of condensate-membrane interaction model. Condensates maintain an intrinsic contact angle (θC) while adopting a double-spherical cap geometry with a deflection angle above (*φ*) and below (*φ*) the GUV membrane plane. The condensate-membrane-solution system is partitioned by condensate-solution (*γ*DS), condensate-membrane (*γ*DM), and membrane-solution (*γ*MS) surface tensions (Nm-1) (yellow arrows). **b,** Representative fluorescence image of condensate-membrane interaction model. The condensate-membrane-solution contact line delimits the above membrane and below membrane spherical caps. **c,** (top) Representative fluorescence images of N protein condensate (cyan) GUV membrane (magenta) interactions in the absence of RNA (left), in the presence of 0.5 nM F3RNA (middle), and in the presence of 0.1 nM vgRNA (right). (bottom) Gradient of interfacial energy (E’(θC,*φ*)) as a function of intrinsic contact angle (θC) and above membrane deflection angle (*φ*). Normalized membrane bending energy (*κ*bend - Nm2) and condensate-membrane-solution line tension (λ - Nm) were determined from the stable solutions (dashed black line) obtained by fitting experimental θC and *φ* measurements (black circles; nno RNA = 4; nF3RNA = 9; nvgRNA = 21). Representative fluorescence images (c.top) are associated with their corresponding experimental data points (c.bottom) (e.g. (i), (ii)). N protein RNA content altered the condensate-membrane interaction geometries observed by fluorescence microscopy. Model fits indicate that membrane bending energy dominates in the no RNA condition (*κ*bend > *γ*DSRD2) leading to condensate deformation rather than membrane bending. In contrast, condensate interfacial energy increasingly dominates in the presence of RNA (F3RNA: *κ*bend < 0.55*γ*DSRD2; vgRNA: *κ*bend << *γ*DSRD2), leading to membrane bending. Intrinsic contact angles (θC) and above membrane deflection angles (*φ*) measurement were performed in ImageJ. Spatial resolution was limited due to the nature of diffraction-limited fluorescence microscopy. All scale bars, 2 µm.

## Discussion

We observed that condensates of N protein alone adhered to the membrane but lacked the interfacial elasticity to maintain their shape, whereas N protein-RNA co-condensates deformed less, resulting in increased membrane bending. These observations suggest a mechanical competition of adhesion between the condensate and the membrane, condensate deformation, and membrane bending. In the case of condensates formed with just the N protein alone, the interfacial elasticity is low such that adhesive forces cause the condensate to deform and flatten out on the membrane, but the elasticity of the membrane prevents it from bending to wrap around the condensate. Although adhesion is sufficient to maintain contact, the condensate flattens rather than inducing membrane curvature, as the membrane’s bending rigidity prevents wrapping. This behavior is reflected in the low aspect ratio of the condensate and the shallow contact angle imposed on the membrane. For N protein condensates formed with 5′ or packaging RNA, their interfacial elasticity is high enough to maintain near spherical shape such that some membrane bending can occur, but the elasticity of the membrane distorts the shape of the condensate to the point that, again, the angle of curvature that the membrane must follow to envelope the condensate is too high to overcome membrane elasticity. Finally, N protein-vgRNA condensates maintain their shape reflecting a balance between opposing elasticities of the membrane and interfacial elasticity of the condensate to allow for its full envelopment. In support of this model, optical trapping measurements show that the presence of RNA causes N protein condensates to become more resistant to deformation.

This study reveals a mechanistic model for SARS-CoV-2 virion maturation, which explains existing results and can be extrapolated to MERS, other coronaviruses that form biomolecular condensate nucleocapsids and it could also be a general mechanism for the selective maturation of RNA virus families with larger genomic RNAs like closterovirthidae. Our data also provides a clue to decoding one of the longstanding questions in the field - how do RNA viruses specifically package vgRNA into virions despite a broad affinity of the N protein to host cellular RNAs. In our model, vgRNA competes out potential cellular RNA-containing nucleocapsids by imparting optimal material properties to the nucleocapsid to drive membrane envelopment. It remains possible to generate virus-like particles in cells simply by expressing the SARS-CoV-2 viral N protein and integral structural proteins E and M^57^. Clustering of integral membrane proteins, such as E and M, on the ERGIC membrane, is expected to reduce membrane stiffness. This reduction may counteract the lower stiffness associated with N protein–cellular RNA condensates, thereby facilitating membrane envelopment. This could also explain why N protein-viral RNA condensates appear to be mutually exclusive from N-M protein condensates in vitro^12^.

It has been suggested that the phosphorylation state of the N protein could govern membrane or N protein-RNA interactions to prevent virion maturation^56^. In our model, phosphorylation could prevent membrane envelopment of the nucleocapsid condensate by reducing either membrane adhesion or condensate elasticity. While post-translational modifications like phosphorylation can affect phase separation^18^, our data show that RNA content of the full-length unmodified N protein condensates is a strong determinant of membrane-based interactions, including bending and encapsulation. The nanoscale hubs observed within the N protein condensates also underscore the dynamic nature of protein-RNA interactions that could affect both viscous and elastic behaviors at a contact interface. Oppositely charged biopolymers are known to form highly dynamic complexes^58^, but the size, sequence and secondary structures formed by viral RNAs could provide a further layer of complexity to the condensate material properties. Moving forward, it would be crucial to advance the methods for developing appropriate thermodynamic models of such dynamic complexes.

Our model provides a new way to think about drug targeting of viral maturation, which will be applicable to other highly pathogenic human coronaviruses like MERS-CoV along with bat coronaviruses like PDF-2180, NeoCoV, and HKU5-CoV that have been identified as high risk for emergence in the human population. Recently, two groups discovered two small molecule inhibitors of SARS-CoV-2 assembly that bind to the membrane protein M and alter its conformational equilibrium^59,60^. M exists in two conformations, a long and a short form, each leading to distinct molecular organization on the membrane and differences in the mechanical stiffness of the virion. The long form results in a more rigid virion, with a narrower range of curvature and spike protein clustering, whereas the short form results in more flexible virions. As discussed above, the M protein may contribute to membrane envelopment of the N protein-vgRNA condensate by modulating membrane elasticity, and it will be interesting to investigate whether the altered conformations induced by the two small molecules lead to membrane stiffening that opposes membrane envelopment. Our findings also provide a framework to delineate new mechanisms of viral genome assembly in other positive and negative-sense strand viruses that show phase separation of IDR-containing RNA-binding capsid proteins^61–66^.

## Methods

### Cloning and expression vectors

The DNA sequence encoding the N protein (full length and C-terminal Cys variant) was synthesized and codon-optimized for *E. coli* expression (Biobasic). The sequence was designed with flanking BamHI and EcoRI restriction sites and cloned into the pGEX-TEV vector^67^, which encodes an N-terminal GST tag followed by a Tobacco Etch Virus (TEV) protease cleavage site upstream of the N protein sequence. The resulting plasmid was transformed into *E. coli Topp2* cells via heat shock and selected on LB agar plates containing 100 mg/mL ampicillin. Plasmid integrity was confirmed by DNA sequencing before transformation into the expression host.

### N protein expression and purification

For protein expression, an overnight *E. coli* culture grown in Luria Broth (LB) medium supplemented with 100 mg/mL ampicillin was diluted into 3 L of LB with antibiotics and incubated at 37°C until reaching an O.D.600 of 0.6–0.8. Protein expression was induced with 0.1mM IsoPropyl-β-D-ThioGalactopyranoside (IPTG; Inalco) for 4 h at 30°C. Cells were harvested by centrifugation and stored at −20°C. Pellets were resuspended in lysis buffer (20 mM Tris-HCl pH 7.5, 1 M NaCl, 5 mM DTT) and lysed using a French Press. The lysate was cleared by centrifugation (1 h, 35,000 rpm, 4°C) and the supernatant incubated with Glutathione Sepharose 4B (GSH; Cytiva) resin for 1 h at 4°C. The resin was washed five times with lysis buffer and subsequently incubated with TEV protease in cleavage buffer (20 mM Tris-HCl pH 7.5, 500 mM NaCl, 5 mM DTT) for 1 h at room temperature. The cleaved N protein was collected and further purified by gel filtration chromatography using a Sepharose 12 10/300GL column (GE Healthcare) in either 20 mM Tris-HCl pH 7.5, 500 mM NaCl, 5 mM DTT or 20 mM Tris-HCl pH 7.5, 1 M NaCl, 5 mM DTT. The wild-type N protein eluted at 12.65 mL in the 500 mM NaCl buffer, while the Cys variant N protein eluted at 13.75 mL in the 1 M NaCl buffer. The purified proteins were stored at −80°C prior to further experimentation. N protein was tagged with Atto 488 NHS ester (Millipore Sigma 41698) or AFdye 488 maleimide (Click Chemistry Tools 1520-1) for the C-terminal Cys variant according to the manufacturer’s protocol.

### Genomic RNA purification

Vero76 (CRL-1587, ATCC) cells were cultured in Dulbecco’s minimum essential medium (DMEM; Gibco, Cat# 11965118) supplemented with 10% FBS, 1% L-Glutamine, and 1% Penicillin/ Streptomycin (Pen/Strep; Gibco, Cat#15-140-122). Calu-3 (HTB-55, ATCC) cells were cultured in DMEM supplemented with 10% FBS and 1% Pen/Step. Vero76 cells were used to propagate SARS-CoV-2 using a previously published protocol^68^. A clinical isolate of ancestral SARS-CoV-2 (SARS-CoV-2/SB2) was used for infection studies following sequence validation using next-generation sequencing^68^. Virus stocks were thawed once and used for an experiment. A fresh vial was used for each experiment to avoid repeated freeze-thaws. All work with infectious SARS-CoV-2 was performed in a containment level 3 (CL3) laboratory at the Vaccine and Infectious Disease Organization (VIDO), University of Saskatchewan using approved protocols. For viral infection, Vero76 or Calu-3 cells were seeded at a density of 3 x 10^5^ cells/well in 6-well plates. When the cells reached 80% confluency, Vero76 and Calu-3 cells were infected with SARS-CoV-2 at an MOI of 0.01 and 0.1, respectively. Infected cells were incubated at 37°C for 1 h with gentle rocking every 15 min. After 1 h, virus inoculum was aspirated, cells were washed with phosphate buffered solution (PBS), and growth media was added on the cells. For Vero76 cells, the supernatant containing SARS-CoV-2 virus particles was harvested at 48 h post infection (hpi) and SARS-CoV-2 genomic RNA was extracted using QIAamp Viral RNA Mini Kit (Qiagen, Cat# 52906) as per manufacturer’s instructions. Cellular RNA from SARS-CoV-2 infected Calu-3 cells was harvested at 48 hpi and RNA was extracted using the RNeasy Plus Mini Kit (Qiagen, Cat# 74134) following manufacturer’s protocol.

### In vitro transcription of viral RNA fragments

A 1kb 5′end region of the vgRNA previously shown to enhance phase separation of N protein invitro^21^, a short packaging signal sequence (1.4kb)^9^, and a size control for the packaging RNA were also synthesized by in vitro transcription and purified. Double-stranded DNA templates encoding 5′, F3 and F3 control RNAs were created by inserting PCR fragments encoding each sequence into a pTZ19R-derived vector containing a T7 promoter site. Prior to transcription, the purified plasmids were fully linearized in 3’ by restriction enzyme digestion. 5′, F3 and F3 control RNAs were transcribed in vitro using T7 RNA polymerase (prepared in house) in 40 mM Tris-HCl pH 8.0, 1 mM spermidine, 0.01% Triton X-100, 25 mM MgCl_2_, 25 mM DTT, 4 mM of each NTP (ATP, CTP, UTP and GTP), and 160 µg/mL of linearized plasmid, and the reaction mixture was incubated at 37°C for 3 h. Next, CaCl_2_ was added at a final concentration of 1 mM along with recombinant DNase I and incubated at 37°C for 2 h to fully digest the DNA template. The enzymes present in the sample were then digested by adding proteinase K at 37°C for 1 h. The transcribed RNA was precipitated at −20°C for 1 h in presence of equal volume 2.5 M LiCl_2_, pelleted by centrifugation, washed with 70% cold EtOH, dried and resuspended to approximately 500 nM in 50 mM Hepes pH 7.6. The purified RNA was labeled at the 3’-end by pCp-Cy3 (Jena Biosciences Cat# NU-1706-CY3) using T4 RNA ligase prepared in house.

### N protein phase separation and GUV preparation

Purified N protein (1-4 µM) was introduced into 100 uL phase separation buffer (20 mM Tris pH 7.4, 40-150 mM NaCl), followed by RNA of varying concentrations directly in the imaging chambers (Ibidi microslides Cat#81816) and imaged immediately by spinning disc confocal microscopy. GUVs were generated by the electroformation method. The lipid mixture contained 1,2-dioleoyl-syn-glycero-3-phosphocholine (DOPC, Avanti Polar Lipids) and 1,2-dioctadecenoyl-sn-glycero-3-Phosphoserine (DOPS, Avanti Polar Lipids,) at a 9:1 molar ratio. Vesicles were fluorescently labelled by addition of 0.1 mol% Texas red dye. Lipid films were deposited onto indium-tin-oxide-coated glass plates (ITO) using glass syringes and 4 mM lipid stocks in chloroform, maintained for 2 h under vacuum and subsequently assembled into a chamber with a 2 mm Teflon spacer. The chamber was held together by binder clips, filled with 170 mM sucrose solution, connected to a function generator (EspoTek Labrador) and a direct current of 3.5 V and 10 Hz frequency was applied for approximately 2 h at room temperature. GUVs were carefully collected and stored at 4°C until use. N protein condensates pre-formed in phase separation buffer were gently mixed with the GUV preparation and imaged immediately.

### Zeta Potential measurement of N protein condensates

Zeta (ζ) potential of N-protein condensates was measured by electrophoretic light scattering (ELS) using a Litesizer DLS 701 instrument (Anton Paar) with continuously monitored phase analysis light scattering (cmPALS). Purified N protein was diluted to a final concentration of 2 µM directly in the cuvette containing phase-separation buffer (20mM Tris pH 7.4, 40mM NaCl). Where indicated, RNA was added at the specified concentrations to induce N-protein–RNA co-condensate formation. Samples were gently mixed without introducing bubbles and allowed to equilibrate for 5 min at room temperature. Measurements were performed in a low-volume Univette reusable cuvette using 5 mV voltage and Smoluchowski correction following the manufacturer’s guidelines and Kalliope software. In short, an electric field was applied to the sample and particle electrophoretic mobility was determined from the Doppler frequency shift of scattered laser light. Each reported value represents the average of 25 technical measurements acquired sequentially from the same sample. Independent experiments were performed in triplicate using freshly prepared samples. Cuvettes were pre-rinsed with filtered buffer, and buffer-only measurements were used for baseline correction. ζ-potential values are reported as mean ± s.d. across independent experiments.

### Optical Tweezer Indentation Assay

N protein-RNA co-condensate indentation assays were performed using the LUMICKS C-Trap system. A 1064 nm 20 W CW trapping laser (IPG Photonics, YLR-20-LP) was used to trap 500 nm diameter PEGylated polystyrene probes. Probes were trapped at 5% overall laser power for all experiments – trapping laser powers were set to 5% and 15% for condensate formed in the absence and presence of RNA respectively resulting in the following average trap stiffnesses: k_trap, vgRNA_= 0.06 pNnm^−1^; k_trap, F3RNA_= 0.04 pNnm^−1^; k_trap, FCRNA_= 0.02 pNnm^−1^; k_trap, noRNA_= 0.02 pNnm^−1^ (Extended data Fig. 9). Using the traps, the probes were steered using motorized piezoelectric mirrors (MadCityLabs, NanoMTA2X) which are conjugate to the back focal plane of the objective lens (60x 1.2NA water immersion, Nikon, Plan apo VC NA1.2). Force detection relies on back-focal plane interferometry. The trap light is collimated by a condenser lens (Leica, P 1.40 OIL S1 11551004) and projected onto a position sensitive detector (PSD, Silicon Sensor International AG, DL100-7PCBA3). The PSD is conjugate to the back focal plane of the objective lens. The PSDs provide voltage signals that reflect the forces exerted on the trapped probe and the probes displacement because of the external forces. Trap force calibration was achieved by fitting the power spectrum of the thermal fluctuations, as measured by the PSD, to a Lorentzian power spectrum. Individual trap calibrations were performed prior to each condensate measurement. Trap calibrations were performed in the system’s dilute phase where the viscosity of the medium was assumed to be that of water at 298 K – 0.89 mPa*s. Using a custom Python code, sequential 0.40 µm linear indentation-retractions cycles were conducted at a constant trap velocity of 1 µm/s. A maximum of 5 indentations were performed per condensate. A custom MATLAB code was implemented to extract condensate FICs. The position of the trapped probe was calculated as *x_probe_ = x_trap_ − F/K_trap_*.

### Condensate-membrane indentation analysis

Dual color fluorescence images of condensate-membrane interactions were analyzed in ImageJ. The fluorescence channels were split to obtain separate images of the condensates and GUV membranes. We manually measured the intrinsic contact angle and the above membrane deflection angles (Fig. 4a-b). We did not consider condensates that had been completely engulfed by GUV membranes. Spatial resolution was limited due to the nature of diffraction-limited fluorescence microscopy.

### FLCS

All FLCS measurements were carried out on a PicoQuant Luminosa single-photon counting confocal microscope (inverted Olympus IX73 body) running on Windows 11 with GPU-accelerated (OpenGL) software control. Excitation was provided by a 485 nm diode laser the power set to approximately [20–50 μW] at the sample. The beam was directed through a 60x, 1.2 NA water-immersion objective (UPLSAPO series). The emitted fluorescence was collected in a parallel detection scheme involving both a single-photon avalanche diode (SPAD) and a hybrid photomultiplier detector (PMA Hybrid-40). To minimize after pulsing contributions from the SPAD, data were analyzed in cross-correlation mode between the two detectors. The system’s time-correlated single-photon counting (TCSPC) unit (MultiHarp 150 8P) features up to 5 ps time resolution and a dead time below 1 ns, enabling precise lifetime measurements. Software based features allowed direct monitoring of excitation power in μW, while the proprietary SymPhoTime 64 software provided integrated FCS and lifetime analysis workflows. Prior to sample measurements, the confocal volume was calibrated using a dilute Atto 488 solution. Fitting the diffusion data yielded an observation volume of approximately 0.3 fL, ensuring accurate count rates and diffusion times for FLCS experiments. Prior to each experiment, the beam path was aligned using the built-in sample-free autoalignment (SFA) feature of the PicoQuant Luminosa, after which a known dye standard (AF488 maleimide) was used to calibrate the confocal volume, ensuring a consistent focal spot size and shape. Typical FCS measurements were then recorded for 60 s per sample spot to achieve reliable photon statistics and autocorrelation curves. Throughout the experiment, laser power at the sample (e.g., 20–50 μW) was monitored and maintained via the CalEx feature, helping to minimize any fluctuations in fluorescence emission. To further reduce photobleaching, illumination was restricted to the actual measurement time, and lower laser powers were employed whenever feasible; additionally, the measurement spot was moved or scanned slightly between acquisitions to avoid prolonged illumination of the same region. Autocorrelation curve fitting was performed using PicoQuant’s OnlineFCS module, which harnesses GPU-accelerated fitting algorithms in real time. This allowed immediate visualization of correlation functions and diffusion parameters, facilitating rapid optimization of acquisition parameters, such as integration time or laser power, based on preliminary results.

### Confocal Line FRAP

Confocal line FRAP measurements were performed on a Abberior STED microscope operated in confocal mode. Condensate samples were prepared as previously described and imaged in Ibidi wells. A 100x oil immersion objective with a 1.4 NA was used for imaging. A 488 nm laser line was used to image N Protein-AF488, and a 561 laser line was used to image vgRNA-Cy3. Imspector software was used to control the following line scan (xt mode) parameters: laser power, pixel dwell time, length of line scan in space, and number of repetitions. To perform a FRAP-like experiment using this confocal microscope, a single condensate was brought into the field of view. A line scan was centered on the condensate and its length spanned the entire condensate. The laser power was set to 100% and 5 thousand repetitions of the line were done. Immediately after, an xyt scan of the entire condensate was taken over time to record the intensity recovery in the condensate. The regional intensity recovery analysis was performed using custom scripts in Matlab 2024b. For FRAP experiments shown in Fig. 2d,e and Extended data Fig. 6a-c, images were acquired with a spinning-disc confocal system (Quorum) using a 100× oil immersion objective and the fluorescence recovery after photobleaching module of Metamorph software. A region of interest of 0.5-1 µm in diameter was set for photobleaching with a 405-nm laser at 100% power for 150 ms. Images were acquired immediately every 2 s for 2 min. Relative fluorescence intensity of the ROI (*F*) through the stacks was measured using the Fiji image processing package^69^. Additionally, recovery in a nearby unbleached fluorescent droplet (control) and a non-fluorescent background region (*F*_b_) were also measured. The photobleaching rate (*r*) was calculated by comparing the fluorescence of the control region before (*F*_c0_) and after (*F*_c_) photobleaching: *r* = *F*_c_/*F*_c0_. The fluorescence intensity of the ROI was normalized with the background, and the photobleaching rate was calculated, *F*_norm_ = (*F* − *F*_b_)/*r*. Finally, the normalized fluorescence intensity was curve fitted with a one-phase exponential equation of GraphPad Prism software.

### Wavelet Spatiotemporal Image Correlation Spectroscopy (STICS)

Imaging time series were acquired on the Luminosa in fluorescence lifetime imaging (FLIM) mode and exported from SymPhoTime 64 as a per-frame 2D photon-count image stack. To emphasize the moving fluorescent *clusters* (and suppress slowly varying background and pixel-scale noise), each frame was band-pass filtered using a Ricker (Laplacian-of-Gaussian) wavelet with a characteristic length scale set to the mean cluster size previously determined by Image Correlation Spectroscopy (ICS). Cluster transport was quantified using spatiotemporal image correlation spectroscopy (STICS)^48^, which measures directed motion by tracking the displacement of the peak of the space–time intensity-fluctuation correlation function. For each subregion (ROI), we computed the normalized spatiotemporal autocorrelation of intensity fluctuations. The generalized space–time correlation function can be approximated by computing the spatial correlation between pairs of frames, as a function of the time lag separating each pair:

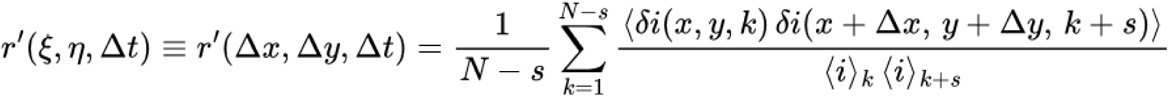

where *(Δx,Δy)* are spatial lags and *τ* is the frame lag. In the presence of coherent drift, the correlation peak shifts approximately linearly with *τ*; the velocity is obtained from the peak displacement. STICS was performed on the wavelet-filtered stack using an ROI size of 16 pixels with an ROI shift of 4 pixels (overlapping tiling), a time-of-interest (TOI) of 5 frames with a TOI shift of 1 frame (sliding temporal window). Local velocity estimates from all ROIs (and time windows) were pooled to generate velocity histograms.

## Supporting information

Supplementary Mathematical modelling

## Mathematical modelling (Supplementary section)

### Data availability

Data supporting the findings of this study are included within the article and its extended data and supplementary figures. Source data are provided with this paper.

## Acknowledgements

We thank P. Garneau for technical assistance in GUV preparation, E. Bajon and N. Stifani for providing expertise with microscopy and C. Roden for critically reading the manuscript. We acknowledge Canadian Institutes of Health Research (CIHR) grant PJT-185976, Human Frontier Science Program grant RGP0034/2017, and the Canada Research Chairs Program (S.W.M.), NSERC (RGPIN-2020-04608) and CIHR(PJT-185997) (A.G.H.), NSERC (RGPIN-2023-03975) and Canada Foundation for Innovation John R. Evans Leaders Fund (CFI-JELF) grant for equipment support (P.W.W), NSERC RGPIN-2023-03837 (JGO), Bristol-Myers-Squibb Chair in Molecular Biology at the Université de Montréal and NSERC RGPIN-2020-05258 (P.L), Canada Research Chairs Program, CFI-JELF and CIHR grants (PPE-192108, PJT-195787) (A.Banerjee). VIDO receives operational funding from the Government of Saskatchewan through Innovation Saskatchewan and the Ministry of Agriculture and from the Canada Foundation for Innovation through the Major Science Initiatives.

## Competing Interests

The authors declare no competing interests.

---

**Extended data Fig. 1:**
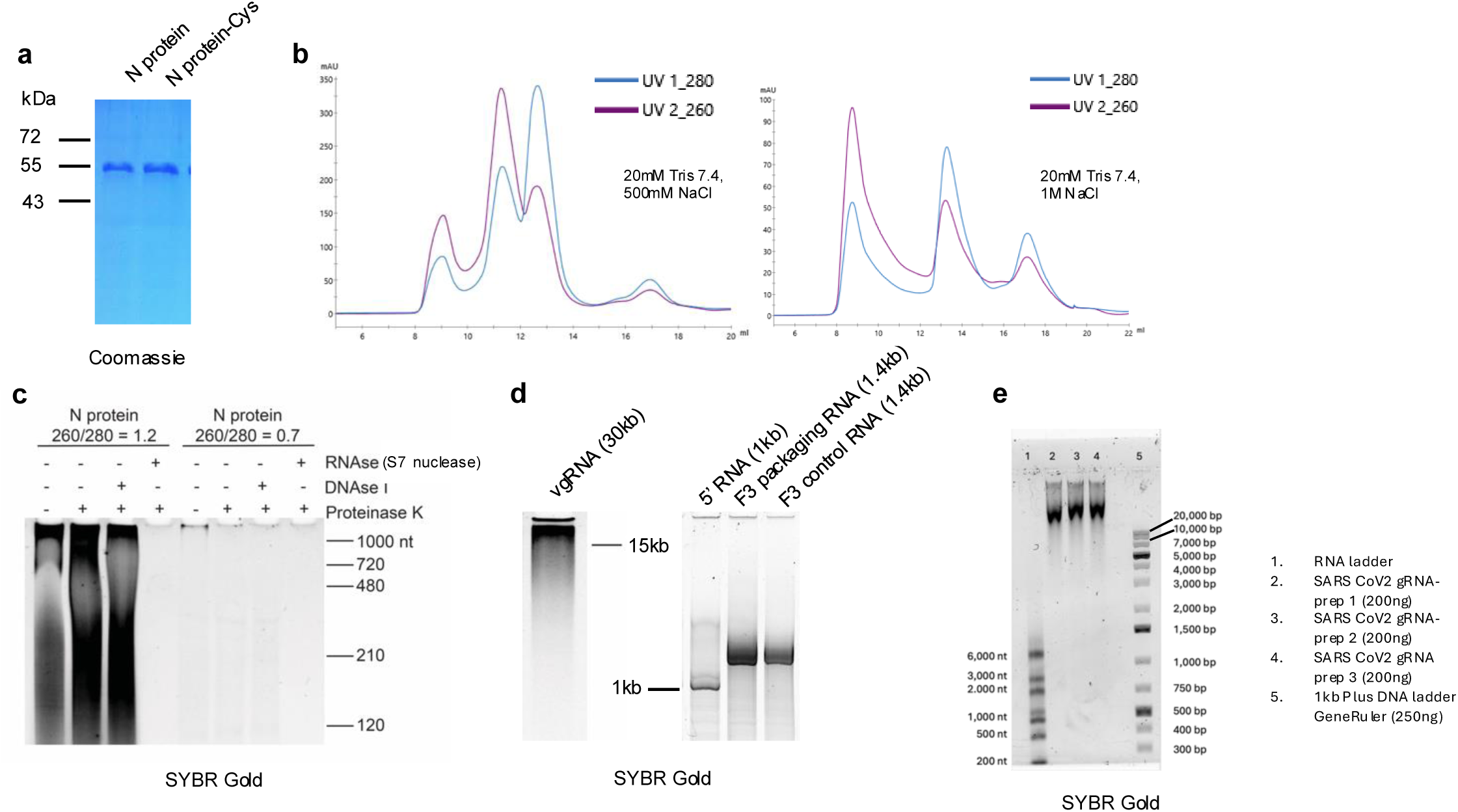
Purified N protein and viral RNA used for phase separation assays: **a,** Coomassie-stained SDS-PAGE gels of 1 µg of the indicated purified proteins. **b**, Chromatogram of N protein purified by gel filtration in a buffer containing 500mM NaCl (left) and 1M NaCl (right). **c,** SYBR gold-stained polyacrylamide gels show RNA as the nucleic acid bound to purified N protein fractions. Protein samples with a high nucleic acid content (260/280=1.2) or a low nucleic acid content (260/280=0.7) were treated with the indicated enzymes prior to loading on polyacrylamide gels. **d**, SYBR gold-stained polyacrylamide gels of purified RNA. **e,** A denaturing agarose showing the purity of vgRNA. 50ng RNA was loaded and visualized using SYBR gold stain.

**Extended data Fig. 2:**
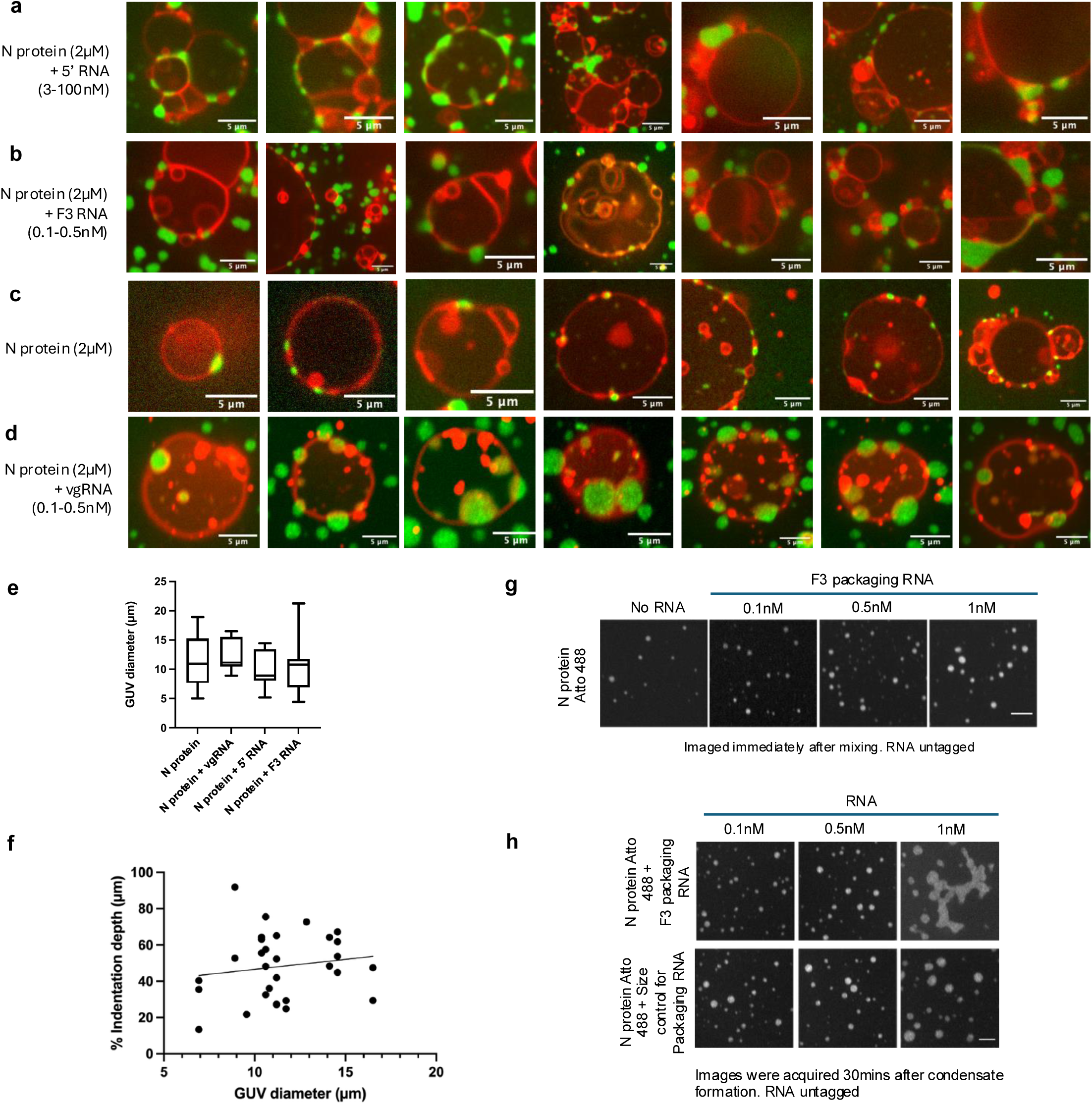
GUV-N protein condensate interaction differs based on RNA content: Confocal images of N-protein condensates (green) interacting with giant unilamellar vesicles (GUVs; Texas Red, in red), in the presence of **a,** 5’ RNA **b,** F3 RNA **c,** no RNA **d,** vgRNA. The merged red-green channel images show the range of interactions with GUVs of variable sizes and condensate diameter. Scale bar 5µm **e,** Quantification of GUV diameter that showed interaction with condensates. **f,** Percent indentation (indentation depth/condensate diameter) plotted against GUV diameter (linear regression: R 2= 0.02345, p= 0.4192; Spearman correlation r= 0.08478, p= 0.6560. **g,** Images of N protein-Atto488 condensates formed in the presence of increasing concentration of F3 packaging RNA. Condensates were imaged immediately after RNA addition. **h,** N protein-F3 RNA condensates (top) imaged 30 mins after RNA addition show aggregate-like structures at 1nM RNA concentration. N protein condensates formed with a size control for the F3 RNA (bottom).

**Extended data Fig. 3:**
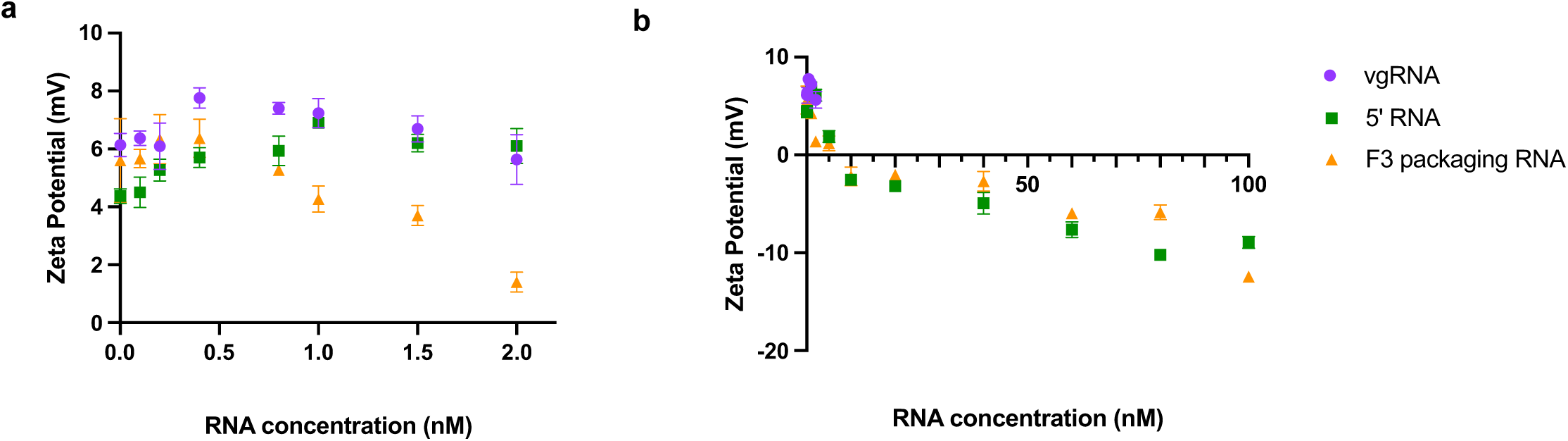
High RNA concentration shifts the Zeta potential of N-protein condensates from positive to negative: **a,** Zeta potential measurements plotted for N protein condensates alone (weakly positively charged, +3 to +5 mV), or with the addition of low RNA concentrations used in GUV experiments. The legend for the RNA is shown to the right, vgRNA (purple), 5’ RNA (green), F3 RNA (orange) **b,** Zeta potential measurements at higher RNA concentrations of 5’ RNA and F3 RNA show a shift to negative values.

**Extended data Fig. 4:**
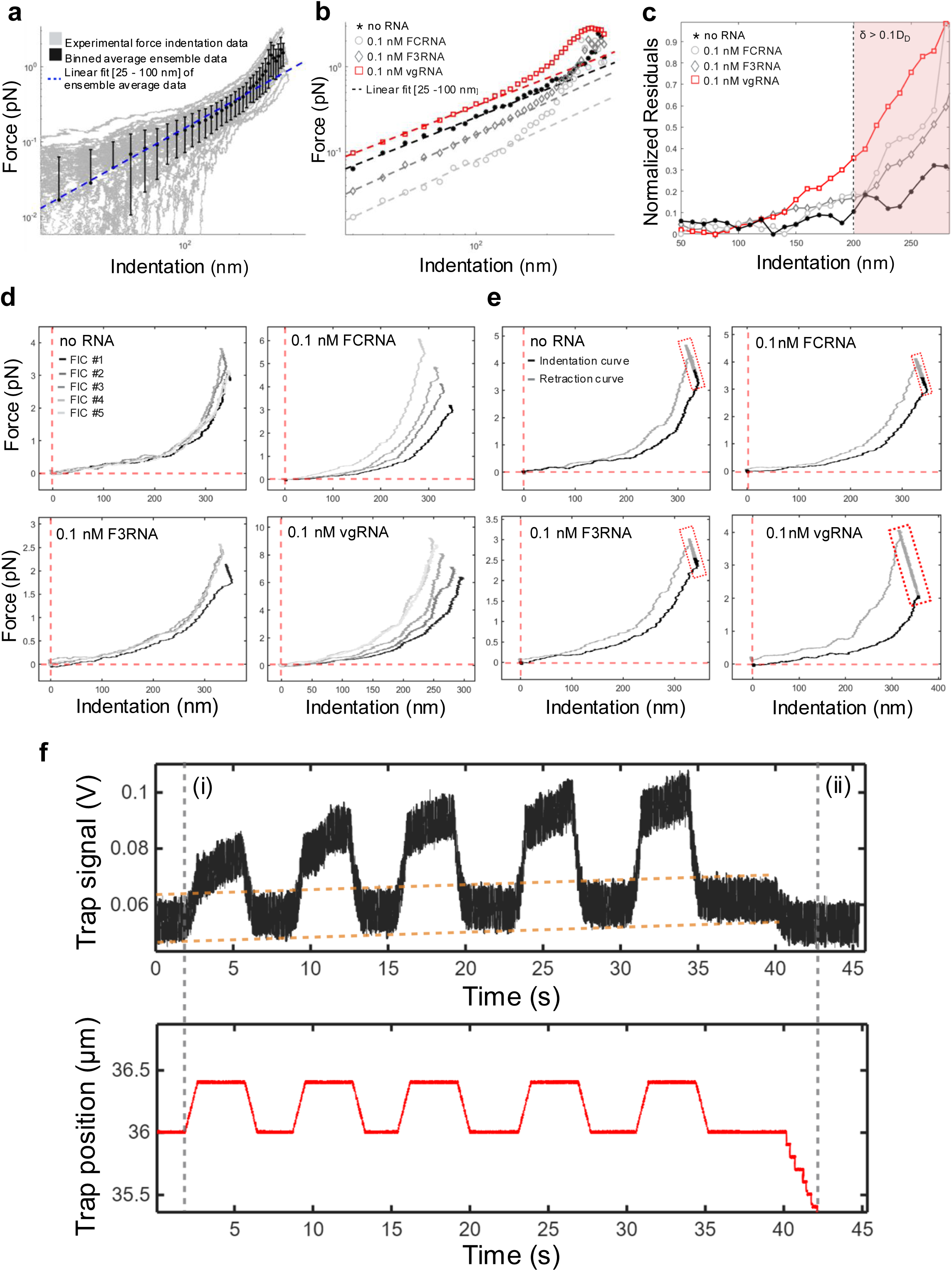
N protein condensates deviate from Hertzian contact mechanics at large indentation depths and after sequential deformations. **a,** All force indentation curves (grey) for one experiment of the vgRNA condition in log-log space. Black symbols indicate the ensemble average and standard deviation. Linear fits to the initial segment (25 – 100 nm) of the ensemble average force indentation curves (blue) indicate deviations from the Hertzian contact model at large indentations (arrow). **b,** Similarly to a, all ensemble average force indentation curves are plotted in log-log space and fitted over their initial segment (25 – 100 nm). **c,** Normalized residuals between the ensemble average force indentation curves and their respective linear fits in b – quantifying the deviation from the linear fit over the course of indentation. In the red region (indentation > 200 nm), all conditions deviate considerably as the small deformation regime is exceeded. Experimental replicates: N_vgRNA_ = 153 FICs; 31 condensates. N_F3RNA_ = 87 FICs; 19 condensates. N_FCRNA_ = 27 FICs; 11 condensates. N_no RNA_ = 19 FICs; 4 condensates. **d,** N protein force indentation curves (FICs) show a characteristic increase upon repeated indentations. Sequential force indentation curves were considered as independent when computing the ensemble average FIC (Fig. 2c) – however, the apparent moduli distribution (Fig. 2d) considers the Hertzian contact fit of each FIC. **e,** N protein condensate indentation-retraction curves show a characteristic force increase when the optical trap is held stationary at its maximum displacement (dashed-red box). The indentation-retraction curves do not suggest that there is observable viscous relaxation. N protein condensates are primarily elastic on the length scales of OT-based indentation assays. Vertical and horizontal red dashed lines represent the prestressed indentations and forces used to zero FICs. **f**, Condensates were prestressed prior to indentation measurements (dashed grey, i). Following sequential condensate indentations, the probe was fully retracted with no observable negative peak, indicating the absence of condensate–probe adhesion (dashed grey, ii). The equilibrium trap signal (dashed orange) and the maximum trap signal increased linearly over the course of sequential condensate indentations, consistent with progressive condensate prestressing resulting from sample drift into the trap.

**Extended data Fig. 5:**
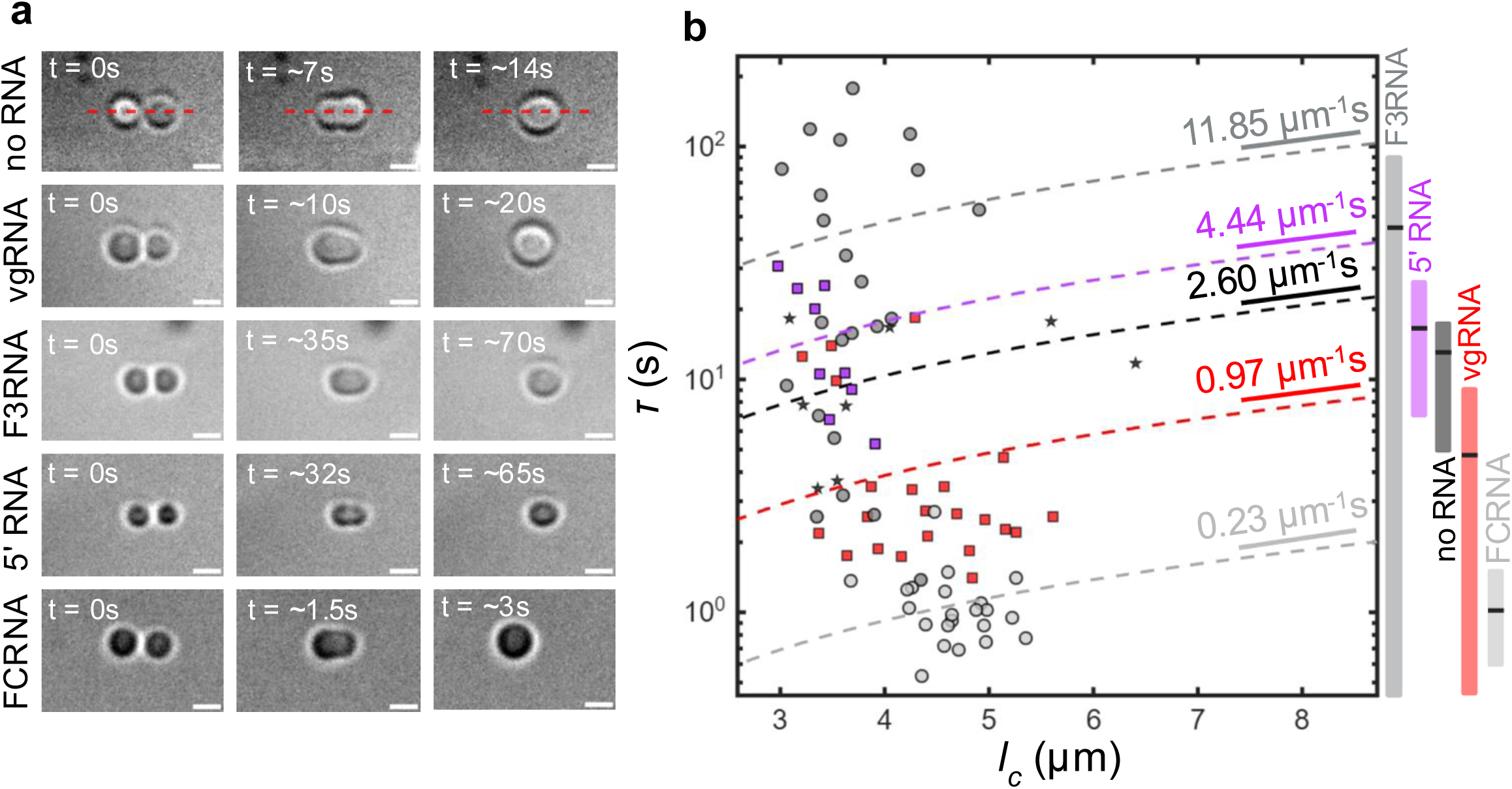
RNA content alters the balance of surface tension elasticity and bulk condensate viscosity. **a,** Representative frames from condensate fusion movies at the onset, during, and end of fusion events. The fusing condensates’ major axis is indicated with a red dashed line. Scale bar: 2μm. Semi-log plot of condensate relaxation times versus initial condensate major axis lengths. RNA conditions’ inverse capillary velocity (dashed lines; µm-1s) were obtained from first-order linear fits of exponential relaxation time constants and major axis lengths measured from time-lapse movies of condensate fusion experiments. Vertical bars indicate the mean ± s.d. of the relaxation time constant for each RNA condition. Sample sizes were: N_noRNA_ = 1 sample, 8 condensates; N_vgRNA_ = 2 samples, 22 condensates; N_F3RNA_ = 2 samples, 23 condensates; N_FCRNA_ = 2 samples, 22 condensates; N_5’RNA_ = 1 sample, 9 condensates.

**Extended data Fig.6:**
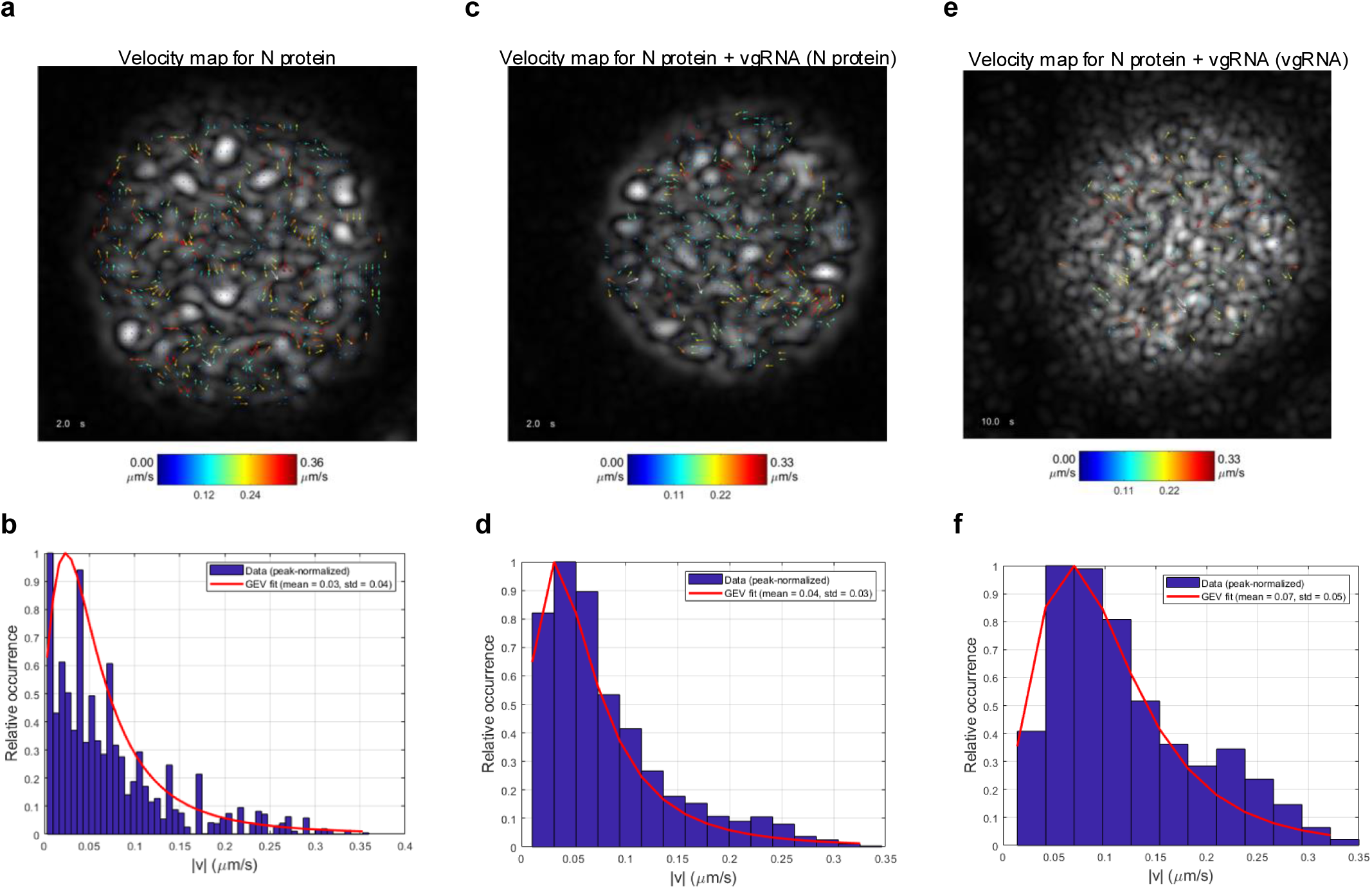
N protein and vgRNA show dynamical behavior on the mesoscale. N protein-AF488 condensates formed without **a,** and in the presence of vgRNA-Cy3 **c,** and **e,**. Imaging time series were analyzed using Richter wavelet filtered spatiotemporal image correlation spectroscopy (STICS) to extract velocity histograms for N Protein in condensates without **b,** vgRNA and with **d,** vgRNA, as well as of **f,** vgRNA-Cy3, of the moving clusters over time. Videos of the time evolution of the velocity are provided in supplementary material. Velocity histograms were fit with a generalized extreme value (GEV) fit, where the parameters µ (mean) and σ (standard deviation) of the GEV are presented.

**Extended data Fig.7:**
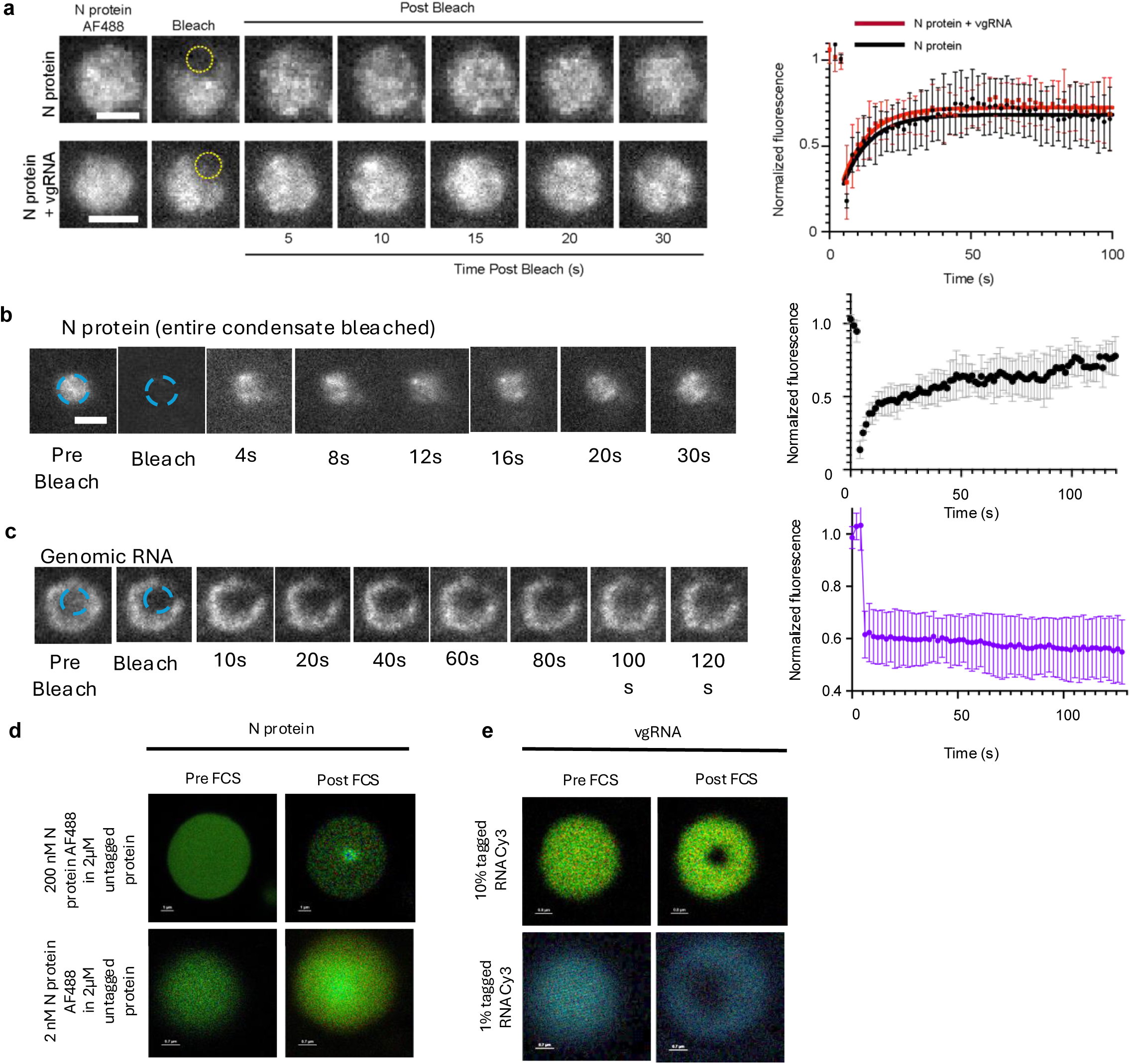
N protein fluorescence recovery begins immediately after photobleaching while RNA remains immobile: **a,** Representative images of N protein-AF488 condensates formed in the presence of vgRNA (bottom) or absence thereof (top) before and after photobleaching. The bleached region of interest is outlined in yellow. Scale bar, 2 µm. Panel on the right shows the quantification of Fluorescence recovery after photobleaching; error bars are mean ± s.e.m. for n = 5 condensates from 3 independent experiments. One-phase exponential equations were fitted to the curves. **b,** FRAP of N protein-AF488 condensates formed in the vgRNA RNA showing N protein recovery or **c,** vgRNA recovery before and after photobleaching. The bleached region of interest is outlined in blue. Scale bar, 2 µm. Quantification of fluorescence recovery is shown on the right. Error bars are mean ± s.e.m. **d,e,** A higher concentration of AF-488 tagged N protein results in an enhanced fluorescence signal during FLCS: Fluorescence lifetime correlation spectroscopy (FLCS) measurements of **d,** N protein-AF488 and **e,** vgRNA-Cy3 taken inside condensates. Images were taken prior FCS and post the 120-second measurement time. The top row shows condensates formed with a higher concentration of labelled N protein or RNA relative to the concentrations used in the bottom row.

**Extended data Fig. 8:**
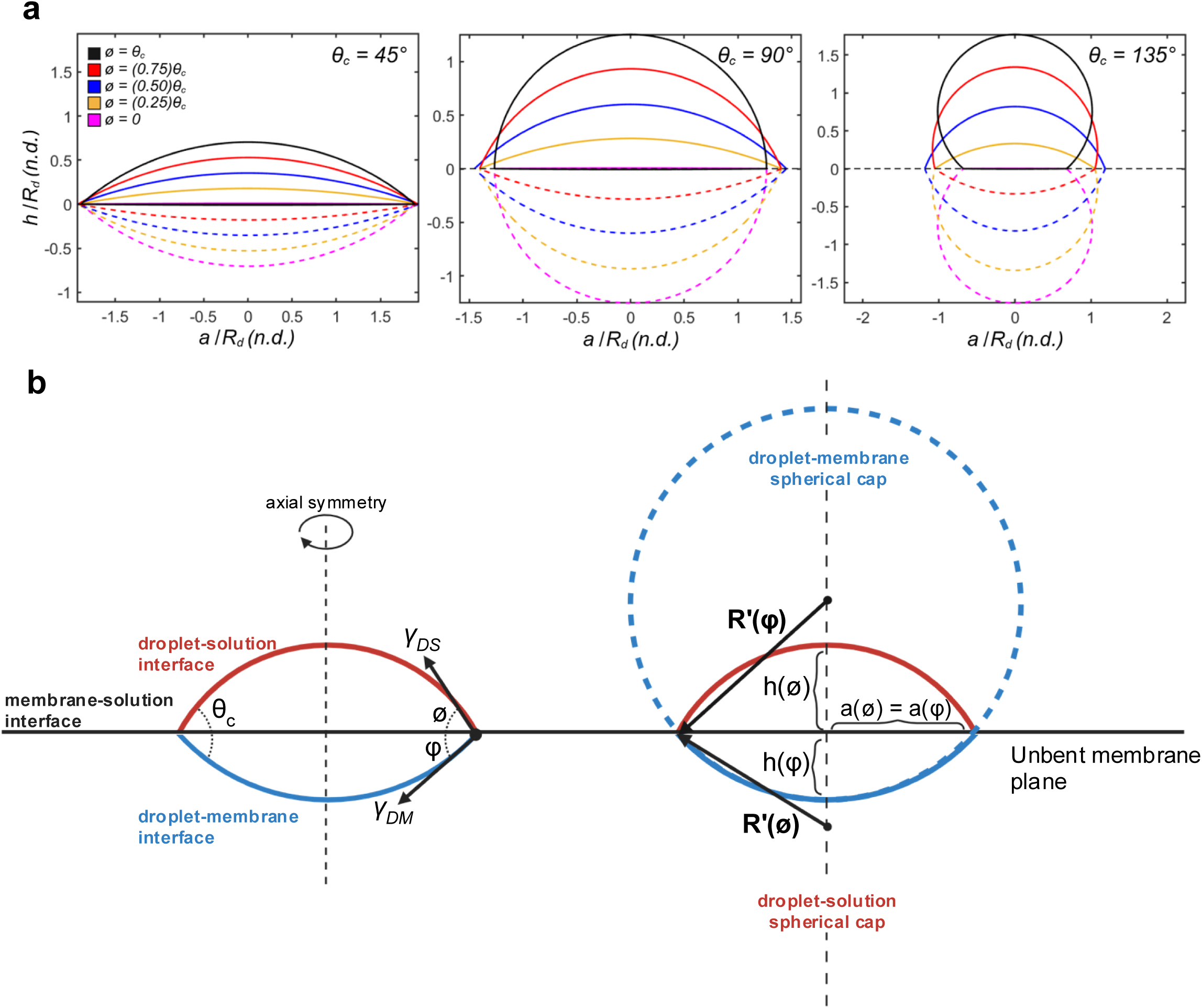
Double spherical cap geometries deflected above and below the membrane plane. **a,** Representative double spherical cap geometries deflected above (solid lines) and below (dashed lines) the membrane plane (h/R_d_ = 0). Left, intrinsic contact angle θ_c_ = 45°; middle, θ_c_ = 90°; right, θ_c_ = 135°. Above-membrane deflection angles (*ϕ*) vary from 0 (pink) to θ_c_ (black). The base radius (a) and height (h) are normalized by the condensate spherical radius (R_d_), assuming conservation of condensate volume during membrane interaction. **b,** (left) Condensates maintain interfaces with the surrounding protein depleted phase (red) and the GUV membrane (blue). Condensate maintain an intrinsic contact angle (*θ_c_*) which is can be decomposed into above membrane and below membrane deflection angles, *ϕ* and *φ* respectively. *ϕ* and *φ* ∈ [0*, θ_c_*] such that *ϕ* + *φ* = *θ_c_*. Condensate-membrane interaction geometries form double-spherical caps that are axially symmetric about their center. (right) Condensate-membrane interactions adopt double-spherical cap geometries composed of an above membrane cap (red) and a below membrane cap (blue). Both spherical caps share a common base and possess their own height. Both spherical caps are also described by spheres of radius *R’(ϕ)* and *R’(φ)* positioned some distance from the unbent membrane plane.

**Extended data Fig. 9:**
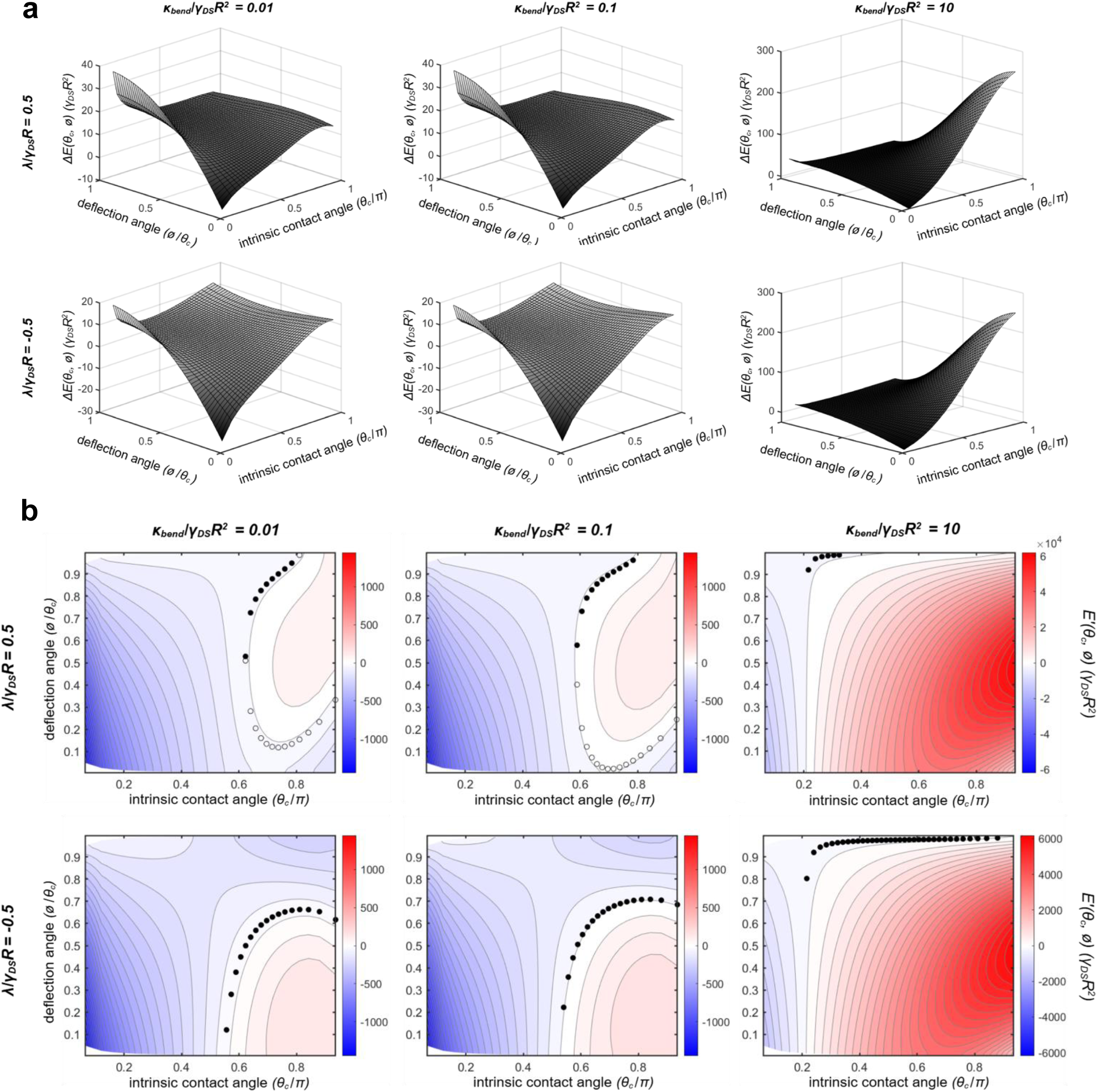
Representative free energy landscapes, *ΔE(θ_c_,ϕ*) = *E(θ_c_,ϕ) - E_0_,* and gradients of condensate-membrane interactions. (left to right) Free energy landscapes and gradients for increasing non-dimensional membrane bending energies (*κ_bend_*/*γ*_DS_R_D_2); 0.01 (left); 0.1 (middle); 10 (right), respectively. (top to bottom) Free energy landscapes for positive (top) and negative (bottom) non-dimensional line tensions (λ/γ_DS_R_D_); 0.5 (top); −0.5 (bottom), respectively. Normalized deflection angles of *ϕ* = 1 correspond to black geometries in Figure S1. Intrinsic contact angles of θ_c_ = π/2 correspond to middle panel geometries in Figure S1.

**Extended data Fig. 10:**
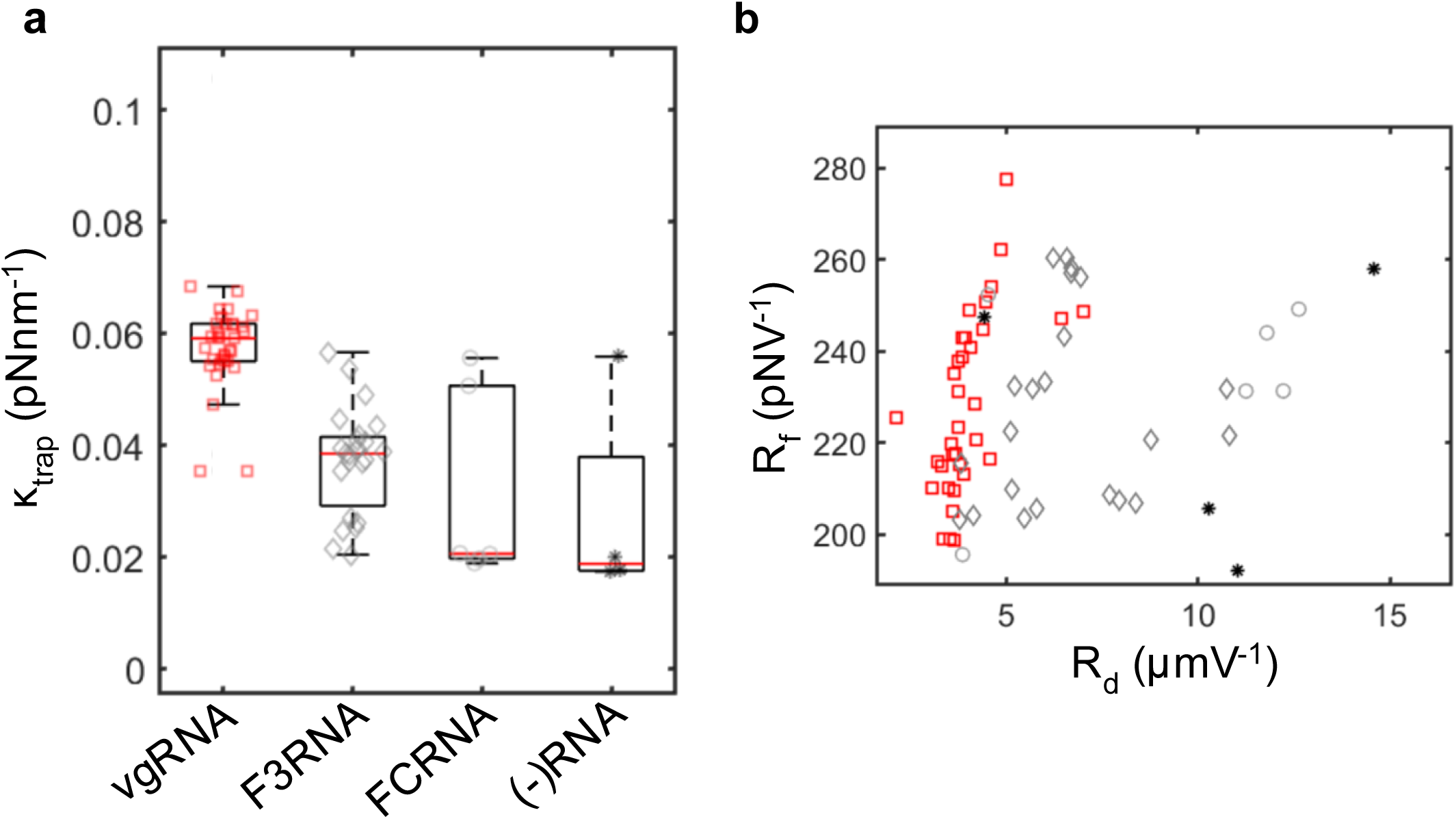
Calibrated optical trapping parameters for all condensate measurements. **a,** k_trap_ (pNnm-1) parameters obtained from fitting the power spectrum of thermal fluctuations to a Lorentzian power spectrum. Individual condensate calibrations are overlaid with their respective RNA conditions. **b,** Scatter plot of optical trap force (R_f_-pNV-1) and displacement (R_d_-µmV^−1^) responses.

## Model of Membrane Bending by Adhesive and Contact Energies

The condensate-membrane interactions can be described using continuum theories of interfacial, contact line, and membrane bending energies (Lipowsky, 2018; Lipowsky *et al.*, 2023; Agudo-Canalejo *et al.*, 2021; Mangiarotti *et al.*, 2023) The free energy of condensate-membrane interactions is given by:

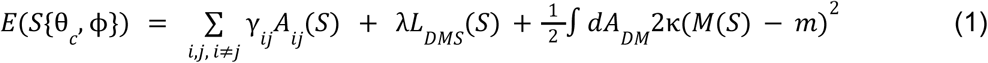

Where *S{θ_c_, φ}* is the shape functional defined by the condensate-membrane intrinsic contact angle (*θ_c_*) and the above membrane deflection angle (*φ*) (Extended data Fig. 8) (Favetta *et al.*, 2024). Intrinsic contact angles can vary from *θ_c_ = 0* in the case of complete membrane wetting (the formation of a thin film or complete condensate engulfment), to *θ_c_ = 180* in the case of no membrane deformations. Above membrane deflection angles range from 0 to *θ_c_*.

*γ*_DS_, *A_DS_*; *γ*_DM_, *A_DM_*; *γ*_MS_, *A_MS_* are the condensate-solvent; condensate-membrane; and membrane-solvent interfacial tensions (Nm^−1^) and areas (m^2^) respectively. Interfacial areas and condensate geometry are explicitly calculated assuming a double spherical cap geometry described by the condensate’s intrinsic contact angle and above membrane deflection angle (Extended data Fig. 9).

λ and *L_DMS_* are the three phase line tension (N) and contact line (m) circumference respectively. The three phase contact line is explicitly calculated assuming a double spherical cap geometry described by the condensate’s intrinsic contact angle and above membrane deflection angle (Extended data Fig. 9).

*κ*, *M*, and *m* are the membrane bending energy (Nm), local membrane curvature (m^−1^), and spontaneous membrane curvature (m^−1^) respectively. Local membrane curvature is explicitly calculated assuming a double spherical cap geometry described by the condensate’s intrinsic contact angle and above membrane deflection angle (Extended data Fig. 8b). While GUVs are spherical and possess a spontaneous curvature, we assume that it is negligible compared to the local curvature induced by condensate-membrane interactions such that *m = 0*.

A portion of N protein-vgRNA condensate interactions result in complete condensate encapsulation (Fig. 1d). Fluorescence microscopy images are consistent with encapsulation via membrane neck scission (Fig. 1d) (Knorr *et al.*, 2015). We fit data assuming stable equilibria prior to complete condensate encapsulation. In this regime, the GUV membrane does not undergo any topological transitions and therefore the Gaussian curvature and bending energies need not be considered (Agudo-Canalejo *et al.*, 2021).

The free energy landscapes of the condensate-membrane system were constructed using a mesh-grid over all possible intrinsic contact angles and above membrane deflection angles. All geometries were normalized by the condensate’s radius (*R_D_*) assuming condensates adopt a spherical geometry in the solvent. This gives rise to a convenient energy normalization scale – *γ*_DS_*R*^2^ – representing the condensate’s interfacial energy prior to the initiation of condensate-membrane interactions (Supplemental Methods - Young Equation). The condensate-membrane shape functional and corresponding free energy were calculated for each node of the mesh-grid and interpolated to produce two dimensional free energy surfaces (Extended data Fig. 9a). Free energy surfaces were numerically analyzed to find equilibrium solutions (Extended data Fig. 9b).

### Double Spherical Cap Geometry

Condensate maintain an intrinsic contact angle (*θ_c_*) which is can be decomposed into above membrane and below membrane deflection angles, *φ* and *φ* respectively. In the special case of undeflected membranes, *φ* = *θ_c_* (Extended data Fig. 8). Condensate-membrane interactions adopt a double-spherical cap geometry defined by their above membrane and below membrane radii, *R’(φ)* and *R’(φ)* respectively. Both spherical caps are axially symmetric about their center (Extended data Fig. 9).

The surface tension vectors, *γ*_DS_ and *γ*_DM_, are tangential to their respective spherical caps at the triple contact line. The spherical cap radii, *R’(φ)* and *R’(φ)*, are normal to their respective spherical caps a the triple contact line (Extended data Fig. 8b). It follows that R’(*φ*) is normal to *γ*_DS_ and *R’(φ)* is normal to *γ*_DM_. The right angle formed between *R’(φ)* and *γ*_DS_ is partitioned by the unbent membrane plane into two angles – the above membrane deflection angle (*φ*) and its complementary angle such that their sum is 90°. We can then show that the angle between the above membrane spherical cap radius and the center line of axial symmetry is *φ*. A similar approach is used to show that the angle between the below membrane spherical cap radius and the center line of axial symmetry is *φ* (Extended data Fig. 8b).

Both spherical caps can be parametrized as a function of their respective deflection angles. The spherical caps share a common base, *a(φ*,*φ)*, which is expressed as:

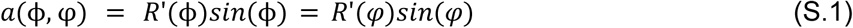

Each spherical cap has its own height, *h(φ)* and *h(φ)*, beginning at the unbent membrane plane and extending to their respective spherical interface. We can express height of the above membrane spherical cap as:

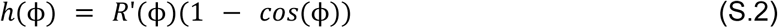

An analogous expression is derived for *h(φ)*.

In the limiting case of a sphere resting atop the unbent membrane plane, we can show that *φ* = 180°, *a(φ*,*φ)* = 0, and *h(φ) = 2R_D_*.

Given the shared base of the double spherical cap geometry, we can express *R’(φ)* in terms of

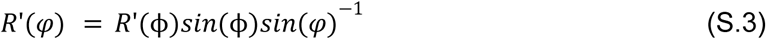

In the limiting case of a sphere resting atop the unbent membrane plane, we can show that *R’(φ)* tends to infinity, indicating that the curvature of the below membrane spherical cap is vanishingly small. To avoid discontinuities, we will use the small angle approximation for *sin(x) ≈ x* for*φ* ∈ [*0, 15*] in our numerical computation of the free energy landscapes.

Assuming that condensate volume remains constant over the timescale of condensate-membrane interactions, we can constrain double spherical cap geometries such that the sum of the above membrane and below membrane spherical caps is equal to the volume of the sphere before condensate-membrane interactions:

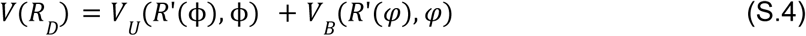

The volume of a spherical cap can be expressed as a function of its base and height:

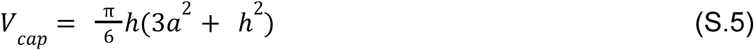

Substituting the expressions for the base and height derived in Eqns. (S.1) and (S.2) into Eqn. (S.5), we express the volume of the spherical cap as a function of its radius and deflection angle:

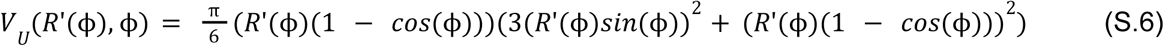

An analogous expression is derived for *V_B_* where the radial terms are substituted with the expression from Eqn. (S.3).

The intrinsic contact angle is maintained across the above and below membrane deflection angles, *φ* + *φ* = *θ_c_*. It follows that *φ* = *θ_c_ - φ*. We can parametrize Eqn. (S.4) as:

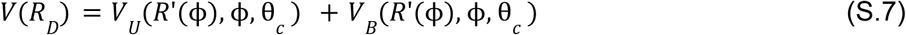

From the volume constraints defined by Eqn. (S.7), we can numerically calculate double spherical cap shape functionals for *θ_c_*∈ [0, 180°] and *φ* ∈ [0*, θ_c_*]. Spherical cap radii are normalized by the initial spherical condensate radius (*R_D_*) (Extended data Fig. 8a).

The interfacial condensate-solvent and condensate-membrane areas are expressed as the area of the above membrane and below membrane spherical caps respectively. The area of the outer shell of a spherical cap is given by:

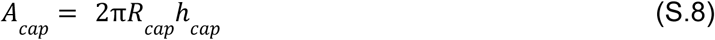

Substituting the expressions for the height derived in Eqn. (S.2) into Eqn. (S.8), we express the condensate-solvent interfacial areas as:

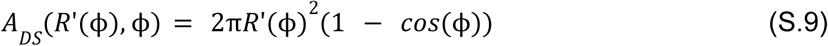

In the limiting case where *φ* = 180°, we find that *A_DS_(R(φ), φ) = 4πR(φ)*^2^. Applying the volume constraints, we recover the surface area of the condensate prior to condensate-membrane interactions, *A_DS_(R(φ), φ) = 4πR*^2^.

An analogous expression is derived for *A_DM_* where the radial terms are substituted with the expression from Eqn. (S.3) and *φ* = *θ_c_ - φ*. In the limiting case of complete condensate engulfment, *φ* = 0° and *φ* = 180°, *A_DM_(R’(φ), φ) = 4πR*^2^.

Prior to condensate-membrane interactions, *A_DM_ = 0* and *A_MS_ =* 4*πR*^2^. As the condensate interacts with the membrane, *A_DM_ = 2πR’(φ)h(φ)* and *A_MS_* = 4*πR*^2^ *-2πR’(φ)h(φ)*. As a result, the total free energy landscape is shifted by the constant interfacial contribution associated with *A_MS_*. Because we are interested only in relative changes in free energy as the condensate–membrane geometry evolves, constant interfacial energy terms are subtracted from the free energy expression.

The three phase contact line, *L_DMS_(S)*, is given by the circumference of the circle with radius *a(φ*,*φ)*.

The local curvature, *M(S)*, is given by the reciprocal of the condensate-membrane spherical cap radius, *R*’(φ)^−1^.

### Young Equation

The condensate-membrane-solvent system is an instance of a solid-liquid-liquid (SLL) system.The GUV membrane is modeled as a deformable elastic solid, whereas the condensate and solvent are modeled as demixed liquid phases. We express the relationship between the three interfacial tensions and the intrinsic contact angle between the condensate and membrane using the Young equation:

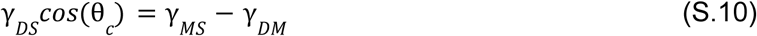

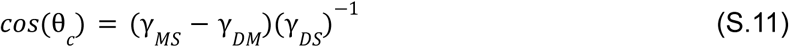

Eqn. (S.11) presents several convenient relationships for the computation of free energy landscapes. We consider all possible *θ_c_*, such that *θ_c_* ∈ [0, 180°]. It follows that *cos(θ_c_)* ∈ [1, −1]. Furthermore, we normalize the free energy landscape by the condensate’s interfacial energy prior to membrane interactions – *γ*_DS_*R_D_*^2^. Eqn. (S.11) provides a physical rational for *γ*_DS_*R_D_*^2^ representing a characteristic energy scale.

## Notes

### Competing Interest Statement

The authors have declared no competing interest.

