## Supplementary Mathematical modelling for "SARS-CoV-2 maturation driven by a mechano-active nucleocapsid-RNA condensate"

### Model of Membrane Bending by Adhesive and Contact Energies.

The condensate-membrane interactions can be described using continuum theories of interfacial, contact line, and membrane bending energies (Lipowsky, 2018; Lipowsky *et al.*, 2023; Agudo-Canalejo *et al.*, 2021; Mangiarotti *et al.*, 2023) The free energy of condensate-membrane interactions is given by:

$$E(S\{\theta_c, \phi\}) = \sum_{i,j, i \neq j} \gamma_{ij} A_{ij}(S) + \lambda L_{DMS}(S) + \frac{1}{2} \int dA_{DM} 2\kappa (M(S) - m)^2 \quad (1)$$

Where $S\{\theta_c, \phi\}$ is the shape functional defined by the condensate-membrane intrinsic contact angle ($\theta_c$) and the above membrane deflection angle ($\phi$) (Extended Data Fig. 8) (Favetta *et al.*, 2024). Intrinsic contact angles can vary from $\theta_c = 0$ in the case of complete membrane wetting (the formation of a thin film or complete condensate engulfment), to $\theta_c = 180$ in the case of no membrane deformations. Above membrane deflection angles range from 0 to $\theta_c$.

$\gamma_{DS}$, $A_{DS}$; $\gamma_{DM}$, $A_{DM}$; $\gamma_{MS}$, $A_{MS}$ are the condensate-solvent; condensate-membrane; and membrane-solvent interfacial tensions ($\text{Nm}^{-1}$) and areas ($\text{m}^2$) respectively. Interfacial areas and condensate geometry are explicitly calculated assuming a double spherical cap geometry described by the condensate's intrinsic contact angle and above membrane deflection angle (Extended Data Fig. 9).

$\lambda$ and $L_{DMS}$ are the three phase line tension (N) and contact line (m) circumference respectively. The three phase contact line is explicitly calculated assuming a double spherical cap geometry described by the condensate's intrinsic contact angle and above membrane deflection angle (Extended Data Fig. 9).

$\kappa$, $M$, and $m$ are the membrane bending energy ($\text{Nm}$), local membrane curvature ($\text{m}^{-1}$), and spontaneous membrane curvature ($\text{m}^{-1}$) respectively. Local membrane curvature is explicitly calculated assuming a double spherical cap geometry described by the condensate's intrinsic contact angle and above membrane deflection angle (Extended Data Fig. 8b). While GUVs are spherical and possess a spontaneous curvature, we assume that it is negligible compared to the local curvature induced by condensate-membrane interactions such that $m = 0$.

A portion of N protein-vgRNA condensate interactions result in complete condensate encapsulation (Fig. 1d). Fluorescence microscopy images are consistent with encapsulation via membrane neck scission (Fig. 1d) (Knorr *et al.*, 2015). We fit data assuming stable equilibria prior to complete condensate encapsulation. In this regime, the GUV membrane does not undergo any topological transitions and therefore the Gaussian curvature and bending energies need not be considered (Agudo-Canalejo *et al.*, 2021).

The free energy landscapes of the condensate-membrane system were constructed using a mesh-grid over all possible intrinsic contact angles and above membrane deflection angles. All geometries were normalized by the condensate's radius ($R_D$) assuming condensates adopt a spherical geometry in the solvent. This gives rise to a convenient energy normalization scale – $\gamma_{DS}R_D^2$ – representing the condensate's interfacial energy prior to the initiation of condensate-membrane interactions (Supplemental Methods - Young Equation). The condensate-membrane shape functional and corresponding free energy were calculated for each node of the mesh-grid and interpolated to produce two dimensional free energy surfaces (Extended Data Fig. 9a). Free energy surfaces were numerically analyzed to find equilibrium solutions (Extended Data Fig. 9b).

#### Double Spherical Cap Geometry

Condensate maintain an intrinsic contact angle ($\theta_c$) which is can be decomposed into above membrane and below membrane deflection angles, $\phi$ and $\varphi$ respectively. In the special case of undeflected membranes, $\phi = \theta_c$ (Extended Data Fig. 8). Condensate-membrane interactions adopt a double-spherical cap geometry defined by their above membrane and below membrane radii, $R'(\phi)$ and $R'(\varphi)$ respectively. Both spherical caps are axially symmetric about their center (Extended Data Fig. 9).

The surface tension vectors, $\gamma_{DS}$ and $\gamma_{DM}$, are tangential to their respective spherical caps at the triple contact line. The spherical cap radii, $R'(\phi)$ and $R'(\varphi)$, are normal to their respective spherical caps at the triple contact line (Extended Data Fig. 8b). It follows that $R'(\phi)$ is normal to $\gamma_{DS}$ and $R'(\varphi)$ is normal to $\gamma_{DM}$. The right angle formed between $R'(\phi)$ and $\gamma_{DS}$ is partitioned by the unbent membrane plane into two angles – the above membrane deflection angle ($\phi$) and its complementary angle such that their sum is $90^\circ$. We can then show that the angle between the above membrane spherical cap radius and the center line of axial symmetry is $\phi$. A similar approach is used to show that the angle between the below membrane spherical cap radius and the center line of axial symmetry is $\varphi$ (Extended Data Fig. 8b).

Both spherical caps can be parametrized as a function of their respective deflection angles. The spherical caps share a common base, $a(\phi, \varphi)$, which is expressed as:

$$a(\phi, \varphi) = R'(\phi)\sin(\phi) = R'(\varphi)\sin(\varphi) \quad (S.1)$$

Each spherical cap has its own height, $h(\phi)$ and $h(\varphi)$, beginning at the unbent membrane plane and extending to their respective spherical interface. We can express height of the above membrane spherical cap as:

$$h(\phi) = R'(\phi)(1 - \cos(\phi)) \quad (S.2)$$

An analogous expression is derived for $h(\varphi)$.

In the limiting case of a sphere resting atop the unbent membrane plane, we can show that $\phi = 180^\circ$, $a(\phi, \varphi) = 0$, and $h(\phi) = 2R_D$.

Given the shared base of the double spherical cap geometry, we can express $R'(\varphi)$ in terms of $R'(\phi)$:

$$R'(\varphi) = R'(\phi)\sin(\phi)\sin(\varphi)^{-1} \quad (\text{S.3})$$

In the limiting case of a sphere resting atop the unbent membrane plane, we can show that $R'(\varphi)$ tends to infinity, indicating that the curvature of the below membrane spherical cap is vanishingly small. To avoid discontinuities, we will use the small angle approximation for $\sin(x) \approx x$ for $\varphi \in [0, 15]$ in our numerical computation of the free energy landscapes.

Assuming that condensate volume remains constant over the timescale of condensate-membrane interactions, we can constrain double spherical cap geometries such that the sum of the above membrane and below membrane spherical caps is equal to the volume of the sphere before condensate-membrane interactions:

$$V(R_D) = V_U(R'(\phi), \phi) + V_B(R'(\varphi), \varphi) \quad (\text{S.4})$$

The volume of a spherical cap can be expressed as a function of its base and height:

$$V_{cap} = \frac{\pi}{6}h(3a^2 + h^2) \quad (\text{S.5})$$

Substituting the expressions for the base and height derived in Eqns. (S.1) and (S.2) into Eqn. (S.5), we express the volume of the spherical cap as a function of its radius and deflection angle:

$$V_U(R'(\phi), \phi) = \frac{\pi}{6}(R'(\phi)(1 - \cos(\phi)))(3(R'(\phi)\sin(\phi))^2 + (R'(\phi)(1 - \cos(\phi)))^2) \quad (\text{S.6})$$

An analogous expression is derived for $V_B$ where the radial terms are substituted with the expression from Eqn. (S.3).

The intrinsic contact angle is maintained across the above and below membrane deflection angles, $\phi + \varphi = \theta_c$. It follows that $\varphi = \theta_c - \phi$. We can parametrize Eqn. (S.4) as:

$$V(R_D) = V_U(R'(\phi), \phi, \theta_c) + V_B(R'(\phi), \phi, \theta_c) \quad (\text{S.7})$$

From the volume constraints defined by Eqn. (S.7), we can numerically calculate double spherical cap shape functionals for $\theta_c \in [0, 180^\circ]$ and $\phi \in [0, \theta_c]$. Spherical cap radii are normalized by the initial spherical condensate radius ($R_D$) (Extended Data Fig. 8a).

The interfacial condensate-solvent and condensate-membrane areas are expressed as the area of the above membrane and below membrane spherical caps respectively. The area of the outer shell of a spherical cap is given by:

$$A_{cap} = 2\pi R_{cap} h_{cap} \quad (\text{S.8})$$

Substituting the expressions for the height derived in Eqn. (S.2) into Eqn. (S.8), we express the condensate-solvent interfacial areas as:

$$A_{DS}(R'(\phi), \phi) = 2\pi R'(\phi)^2 (1 - \cos(\phi)) \quad (\text{S.9})$$

In the limiting case where $\phi = 180^\circ$, we find that $A_{DS}(R(\phi), \phi) = 4\pi R(\phi)^2$. Applying the volume constraints, we recover the surface area of the condensate prior to condensate-membrane interactions, $A_{DS}(R(\phi), \phi) = 4\pi R_D^2$.

An analogous expression is derived for $A_{DM}$ where the radial terms are substituted with the expression from Eqn. (S.3) and $\varphi = \theta_c - \phi$. In the limiting case of complete condensate engulfment, $\phi = 0^\circ$ and $\varphi = 180^\circ$, $A_{DM}(R'(\varphi), \varphi) = 4\pi R_D^2$.

Prior to condensate-membrane interactions, $A_{DM} = 0$ and $A_{MS} = 4\pi R_{GUV}^2$. As the condensate interacts with the membrane, $A_{DM} = 2\pi R'(\varphi)h(\varphi)$ and $A_{MS} = 4\pi R_{GUV}^2 - 2\pi R'(\varphi)h(\varphi)$. As a result, the total free energy landscape is shifted by the constant interfacial contribution associated with $A_{MS}$. Because we are interested only in relative changes in free energy as the condensate-membrane geometry evolves, constant interfacial energy terms are subtracted from the free energy expression.

The three phase contact line, $L_{DMS}(S)$, is given by the circumference of the circle with radius $a(\phi, \varphi)$.

The local curvature, $M(S)$, is given by the reciprocal of the condensate-membrane spherical cap radius, $R'(\varphi)^{-1}$.

### Young Equation

The condensate-membrane-solvent system is an instance of a solid-liquid-liquid (SLL) system. The GUV membrane is modeled as a deformable elastic solid, whereas the condensate and solvent are modeled as demixed liquid phases. We express the relationship between the three interfacial tensions and the intrinsic contact angle between the condensate and membrane using the Young equation:

$$\gamma_{DS} \cos(\theta_c) = \gamma_{MS} - \gamma_{DM} \quad (\text{S.10})$$

$$\cos(\theta_c) = (\gamma_{MS} - \gamma_{DM})(\gamma_{DS})^{-1} \quad (\text{S.11})$$

Eqn. (S.11) presents several convenient relationships for the computation of free energy landscapes. We consider all possible $\theta_c$, such that $\theta_c \in [0, 180^\circ]$. It follows that $\cos(\theta_c) \in [-1, 1]$. Furthermore, we normalize the free energy landscape by the condensate's interfacial energy prior to membrane interactions $-\gamma_{DS}R_D^2$. Eqn. (S.11) provides a physical rationale for $\gamma_{DS}R_D^2$ representing a characteristic energy scale.
